# Neurogranin enhances spontaneous activity and neuronal survival of hippocampal neurons

**DOI:** 10.64898/2026.01.27.701932

**Authors:** Elena Martínez-Blanco, Raquel de Andrés, Esperanza López-Merino, Jose Antonio Esteban, F. Javier Díez-Guerra

## Abstract

Neurogranin (Ng) is a postsynaptic protein highly enriched in forebrain neurons and implicated in synaptic plasticity through its ability to bind calmodulin. However, its impact on neuronal development, network dynamics, and cellular homeostasis remains incompletely understood. In this study, we examined the effects of manipulating Ng expression in primary hippocampal neurons using viral gene delivery, with emphasis on structural, functional, and molecular outcomes. Restoring Ng expression to adult physiological levels enhanced dendritic growth, increased synaptic number, and induced a proximal shift of the axon initial segment, consistent with adaptive responses to increased connectivity. Functionally, Ng markedly increased spontaneous neuronal activity and network synchronization, even under culture conditions that normally show minimal baseline activity. Electrophysiological recordings revealed enhanced burst firing and spike synchrony, indicating strengthened functional coupling rather than increased membrane excitability. Ng-dependent activity required action potential firing and glutamatergic transmission. At the molecular level, Ng increased total calmodulin levels in a binding-dependent manner, reduced overall calcium/calmodulin-dependent protein kinase II abundance while enhancing its relative autophosphorylation, and selectively decreased both total and surface levels of ionotropic glutamate receptors. These changes are consistent with a coordinated homeostatic reorganization of calcium-dependent signaling. Despite robust increases in activity, Ng expression improved neuronal viability, reduced cellular stress markers, and increased expression of the anti-apoptotic protein Bcl-2. Active caspase-3 was selectively elevated without triggering apoptosis, suggesting a non-apoptotic role in activity-dependent structural remodeling. Together, these findings identify Ng as a homeostatic regulator that promotes coordinated network activity, adaptive synaptic remodeling, and neuronal survival.

## Introduction

Neurogranin (Ng), a small protein abundantly expressed in principal neurons of the forebrain [1, 2], is predominantly localized to the somatodendritic compartment, with a notable enrichment in dendritic spines [3]. Ng binds calmodulin (CaM) with higher affinity under low intracellular calcium ion (Ca^2+^) concentrations through its central IQ motif, which adopts a stabilized α-helical conformation upon CaM binding [4]. Among neuronal CaM-binding proteins, Ng is one of the most abundantly expressed in the rodent forebrain [5]. Ng is a substrate of protein kinase C (PKC) [6, 7], which phosphorylates serine 36 within the IQ motif. This phosphorylation disrupts Ng-CaM interaction, thereby regulating the availability of free CaM for downstream signaling. Under resting conditions, Ng limits Ca^2+^/CaM signaling due to its high buffering capacity and because Ng binding reduces CaM’s affinity for Ca² [8]. Elevation of intracellular Ca² levels weakens the Ng-CaM interaction, thereby releasing free CaM.

Ng is a sensitive indicator of synaptic dysfunction and cognitive decline [9]. It has also emerged as an early biomarker for mild cognitive impairment (MCI) and prodromal Alzheimer’s disease (AD) [10], through its detection in cerebrospinal fluid (CSF) [11–14]. Synaptic dysfunction and loss is recognized as one of the earliest pathological events in neurodegenerative disorders, often preceding neuronal death [15]. Both human and animal studies support a link between Ng expression and cognitive performance [14, 16, 17]. Reduced levels of brain Ng have been correlated with cognitive deficits in several contexts, including aging and specific pathological conditions [14]. For example, developmental hypothyroidism [18] and exposure to chemicals such as bisphenol [19], known to reduce circulating triiodothyronine (T3) levels, have been shown to decrease Ng expression and impair cognitive function.

Despite extensive research, the precise role of Ng in synaptic plasticity remains incompletely understood. Studies using Ng knockout (Ng KO) mice have yielded conflicting results: one reported impaired long-term potentiation (LTP) following high-frequency stimulation [20], while another found enhanced LTP alongside deficits in long-term depression (LTD) [21]. Given the complexity of Ca² /CaM signaling, several computational models have been developed to further explore Ng’s function [22–24]. One model [22] predicts that Ng deficiency impairs LTP induced by a single 100 Hz, 1 sec tetanus, suggesting that LTP relies on CaM stored in spines as rapidly dissociating CaM-Ng complexes. Another model [23] proposes a Ca^2+^-dependent dual role for Ng: at moderate local Ca² levels (<10 μM) Ng would suppress Ca² /CaM target activation, whereas at higher Ca² levels Ng would be functionally irrelevant. A third study [24] concluded that Ng reduces the likelihood of CaMKII autophosphorylation. Notably, these models did not consider the effects of Ng phosphorylation by PKC, a process itself influenced by intracellular Ca^2+^ levels.

Many experimental investigations of Ng function have utilized hippocampal organotypic cultures or acute slices. In Ng knockout mice, reduced hippocampal Ng levels are associated with impaired spatial learning [25], diminished LTP, enhanced LTD, and attenuated Ca² transients in CA1 pyramidal neurons following tetanic stimulation [5]. Additional work using organotypic slices has shown that Ng enhances postsynaptic sensitivity and synaptic strength in an activity- and NMDA receptor-dependent manner [26], potentially by targeting CaM within dendritic spines [27]. However, the molecular mechanisms underlying Ng enrichment in active spines remains poorly understood. Previous work identified phosphatidic acid (PA) as a potential Ng binding partner [28], suggesting a possible anchoring mechanism, though this hypothesis remains to be experimentally validated. In vitro studies have also shown that Ng can attenuate the activation of Ca² /CaM-dependent targets [29]. Consistent with this, it has been proposed that small IQ-motif proteins like Ng may serve to constrain the activation or sustained activity of Ca² /CaM-dependent enzymes [30].

Given its unique characteristics, Ng emerges as a promising target for preventive or therapeutic strategies aimed at counteracting cognitive decline. Ng is expressed relatively late in development, displays a highly specific distribution within the forebrain, and -despite its absence not resulting in major anatomical or physiological abnormalities-its deficiency leads mostly to marked cognitive impairments. These features highlight the importance of further elucidating Ng’s functional role in the neural environment. To address this, we conducted an in-depth investigation using primary cultures of embryonic rat hippocampal neurons -an experimental model that offers greater control over environmental variables than in vivo systems or organotypic slice cultures. In our previous work, we found that unlike other synaptic proteins, Ng expression in mature cultured hippocampal neurons does not reach the levels observed in intact brain tissue [31]. We leveraged this finding by comparing unmodified neuronal cultures with those transduced with adeno-associated viruses (AAVs) engineered to drive Ng expression under the control of the excitatory neuron-specific CaMKIIα promoter, enabling extensive and targeted expression of Ng at physiologically relevant levels. Through a combination of biochemical, imaging, and electrophysiological techniques, we found that Ng expression enhances neuronal connectivity, activity, and survival. These results support the hypothesis that Ng acts as a homeostatic regulator of neural network activity. Moreover, its presence appears to mitigate the harmful effects of intracellular calcium surges, which can disrupt metabolic stability and compromise neuronal viability. In summary, our findings reinforce the view that Ng contributes to the functional integrity and resilience of neural networks. Its ability to promote synaptic activity and neuronal survival makes it a particularly attractive candidate for interventions aimed at preserving cognitive function.

## Materials and Methods

### Animals and ethics compliance

Wistar rats were bred at the animal facility of the Universidad Autónoma de Madrid (UAM). All procedures conducted during the study strictly adhered to the Spanish Royal Decree 1201/2005, which governs the protection of animals used in scientific research, as well as the European Union Directive 2010/63/EU concerning the welfare of animals in scientific contexts. Experimental protocols were approved by the corresponding institutional and regional ethics committees, ensuring that the highest standards of animal care and ethical compliance were maintained throughout the research.

### Reagents

Fetal bovine serum (FBS), Dulbecco’s modified Eagle’s medium (DMEM), 0.25% trypsin, Neurobasal media (NB) and B27 supplement were from Thermofisher Scientific. The protease inhibitor cocktail was from Biotools (B14001). Total protein was measured using the Bradford Protein Assay kit (Bio-Rad). Pre-stained protein markers VI (10-245 kDa) were from PanReac-AppliChem. Immobilon-P membranes and ECL western blotting reagents were from Millipore. 1-beta-arabino-furanosylcytosine (AraC) was from Calbiochem (251010). AP5 was from Tocris (ref. 0106), 2-APB and Poly-L-lysine hydrobromide (PLL) from Sigma-Aldrich (refs. 100065 and P2636, respectively), NBQX was from Tocris (ref. 0373), TTX from Alomone Labs (ref. T-500) and MPEP from MedChemExpress (ref. HY-14609A). Paraformaldehyde (PFA) was from Merck. Antibodies and plasmids used in this study are listed in Supplementary Tables S1-S3.

### Primary cultures of rat hippocampal neurons

Primary hippocampal neurons were prepared from embryonic day 19 (E19) Wistar rat embryos as previously described [32]. Embryos were collected and maintained in chilled Hank’s balanced salt solution (HBSS) and hippocampi were carefully dissected free of meninges. Tissue was washed five times in HBSS and incubated with 0.25% trypsin for 15 minutes at 37 °C to facilitate cell dissociation. After trypsin removal by two additional washes in HBSS, tissue was transferred to HBSS containing 1.26 mM CaCl_2_, 0.81 mM MgSO_4_ and 0.04 mg/mL DNase I, and dissociated mechanically using Pasteur pipettes and 22G needles. The resulting cell suspension was filtered through a 70 µm nylon mesh, centrifuged at 300 xg for 5 minutes, resuspended in plating medium (10% FBS/DMEM, 1 mM Pyruvate, 10 mM D-glucose), and counted. Cells were plated at a density of 25,000 cells/cm^2^ onto culture dishes pre-coated with 0.1 mg/mL PLL in borate buffer (pH 8.0), or at 12,000 or 20,000 cells/cm^2^ on 18-mm or 25-mm coverslips, respectively, pre-treated with 0.25 mg/mL PLL. Coverslips were previously sterilized with 65% nitric acid for 24–72 h, extensively washed with double-distilled H2O (ddH2O) and baked at 180 °C. 3 hours after plating to allow cell adhesion to the substrate, the plating medium was replaced with Neurobasal medium (NB) supplemented with B27 and GlutaMAX (Thermofisher). To suppress glial proliferation, 1 µM AraC was added on day in vitro 3 (DIV3). On DIV7, 50% of the culture medium was replaced with fresh NB/B27. Cultures were maintained at 37 °C in a humidified atmosphere with 5% CO and used for experiments during the third week in vitro. All treatments were performed within the thermostatized CO_2_ incubator.

### Preparation of lentiviral and adeno-associated viral particles

HEK-293 T cells were cultured in DMEM supplemented with 10% FBS (Gibco) and maintained at 37°C with 5% CO_2_. Passaging was performed twice a week.

#### Lentiviral particles (LVs) production

All lentiviral constructs are derived from the vector pLOX-Syn-DsRed-Syn-GFP [33], kindly donated by Dr. FG Scholl. HEK-293T cells (3 × 10^6^) were seeded in p100 dishes and transfected 24 hours later with 8 µg of the lentiviral plasmid of interest, 4 µg of pCMVδR8.74, and 2 µg of pMD2.G using PEI MAX (DNA:PEI ratio 1:2). DNA and PEI were mixed in OptiMEM medium (Gibco), incubated for 20 minutes at room temperature (RT), and added dropwise to the cells. After 5 hours of incubation at 37 °C and 5% CO, the transfection medium was replaced with Neurobasal medium. After 48 h, the medium containing lentiviral particles was collected, filtered through a 0.45 µm filter, aliquoted, and stored at -80 °C. Cultures of hippocampal neurons were infected at day in vitro 4 (DIV4).

#### Adeno-associated virus (AAV) production

AAVs were prepared as previously described [34]. HEK-293T cells (7 × 10 ) were seeded in p150 dishes and cultured in 10% FBS/DMEM until 70% confluency, at which point the medium was replaced with 5% FBS/DMEM. Cells were then transfected using the calcium phosphate method with the following plasmids: 12.5 µg of the desired pAAV plasmid, 25 µg of pFδ6, 6.25 µg of pH21, and 6.25 µg of pRV1. Plasmids were diluted and adjusted to a volume of 900 µL using CaCl_2_ (final concentration 125 µM). An equal volume of 2× HBS (274 mM NaCl, 10 mM KCl, 1.4 mM Na_2_HPO_4_, 15 mM D-Glucose and 42 mM HEPES buffer, pH 7.05) was added, the solution mix bubbled and incubated for 20 minutes at RT, then added dropwise to the cultures. After 16 hours of incubation (37 °C, 5% CO ), the medium was replaced by 10% FBS/DMEM. 48 hours post-transfection, cells were washed with PBS and lysed in extraction buffer (150 mM NaCl, 20 mM Tris-HCl pH 8, 0.4% sodium deoxycholate, 50 U/mL benzonuclease (Millipore)). Lysates were centrifuged at 3,000 × g for 15 minutes, and the supernatant applied to a heparin Sepharose column (HiTrap, Cytiva). Eluted fractions were concentrated using Amicon Ultra-4 100 kDa filters (Millipore), filtered through 0.13 µm filters, aliquoted, and stored at −80 °C. Hippocampal neurons were typically infected at DIV7 with AAV particles diluted in half the culture volume of the dish. After an 8-hour incubation, the viral medium was replaced with a 1:1 mixture of the previously collected medium and fresh NB supplemented with B27 and GlutaMAX. To silence Ng expression, the plasmid shNg pA_RC3J1_CAGW (Addgene #92155), encoding an shRNA targeting the sequence gtgacaagacttccctactgt, was used. Prior to shNg AAV production, the GFP coding region of the original plasmid was replaced by mRuby2. As a control, a non-targeting scrambled shRNA (gtgccaagacgggtagtca, Addgene #181875) was used. To preserve neuronal viability, AAV infections for knockdown experiments were performed at DIV10.

### Protein extraction and Western blots

Cells were lysed in extraction buffer containing 50 mM NaCl, 0.5% Triton X-100, 1 mM EDTA, 2 mM DTT, 25 mM Tris-HCl (pH 6.8), and protease & phosphatase inhibitors (Biotool). Lysates were homogenized by 20 passes through a 23G needle and centrifuged at 17,500 × g for 15 minutes at 4 °C. Protein concentrations in the supernatants were determined using the Bradford assay (Bio-Rad) and 15– 25 μg from hippocampal neuron extracts were loaded per well. Proteins were separated by SDS-PAGE on 6%, 10% or 13% polyacrylamide gels (Mini Vertical Protein Electrophoresis System, Cleaver Scientific) under reducing conditions at 120 V for ∼2.5 hours. Protein in the gels was transferred onto PVDF membranes (Millipore) using a semi-dry transfer system (Nyx Technik) at a constant current of 400 mA for 30 minutes in transfer buffer (22.5 mM Tris, 170 mM glycine, 20% methanol). Membranes were blocked with 5% (w/v) skimmed milk in TBS for 1 hour at room temperature with agitation and incubated overnight at 8 °C with primary antibodies in TBS with 0.05% Tween-20. HRP-conjugated secondary antibodies (Jackson ImmunoResearch, 1:15.000) were used for detection with an enhanced chemiluminescence system (ECL, Millipore). Signal acquisition was performed using an Amersham Imager 680 (GE Healthcare Life Sciences), and densitometric analysis was carried out with the open source software Fiji/ImageJ [35, 36].

### Cell surface biotinylation

Hippocampal neurons at DIV16, previously seeded in 6-well plates, were washed with ice-cold PBS and cell surface proteins were labeled for 30 minutes at 4°C by incubation in a 1 ml solution containing the non-permeable Sulfo-NHS-SS-Biotin (1 mg/ml in PBS). The biotinylation reagent was subsequently quenched twice with 100 mM L-Lysine for 30 minutes at 4°C. After three additional washes with PBS, cells were lysed for 30 minutes in lysis buffer (150 mM NaCl, 1 mM EDTA, 50 mM Tris-HCl (pH 7.4), 1% Triton X-100 and proteases inhibitors) and the lysate clarified by centrifugation at 17,500g for 15 minutes. Biotinylated proteins were recovered by incubating the cleared lysate with streptavidin-agarose beads for 90 minutes at room temperature. After washing the beads three times with 1 ml of the lysis buffer, bound proteins were eluted in 2× Laemmli sample buffer at 95°C, separated by SDS-PAGE, and analyzed in Western blot.

### Immunofluorescence

Primary hippocampal neurons (DIV16–17) grown on 18 mm round coverslips were quickly washed with PBS and fixed with 4% PFA in PBS for 20 minutes at room temperature. The fixative was removed, and the cells incubated with 0.2 M glycine (pH 8) for 5 minutes at RT to quench residual PFA. After three PBS washes, cells were permeabilized and blocked for 30 minutes in blocking buffer containing 0.1% Triton X-100, 1% bovine serum albumin (BSA), and 1% heat-inactivated horse serum in PBS. Incubation with primary antibodies (Supplementary Table S3) was conducted overnight at 8°C in PBS supplemented with 1% BSA and 1% horse serum. After three 5-minute PBS washes, secondary antibodies were applied for 1 hour at room temperature in the same buffer. Nuclei were counterstained with DAPI (0.2 μg/mL in PBS) for 5 minutes, followed by washes in distilled water and 96% ethanol. Coverslips were air-dried and mounted using Mowiol. Images were acquired on a Zeiss AxioObsever7 fluorescence microscope and a Hamamatsu Orca Fusion Camera. Excitatory and inhibitory synaptic contacts were identified by immunostaining with antibodies against specific presynaptic and postsynaptic markers (Fig.1c). MAP2 labeling was used to define dendritic compartments and to restrict synapse analysis to regions in close apposition to dendrites. Synaptic contacts were analyzed and quantified using the SynapTrack [37] macro language script implemented in Fiji/ImageJ [35, 36].

### Measurement of axon initial segment length (AIS) length and distance from the soma

Hippocampal neurons were processed for immunofluorescence using antibodies against Ankyrin G (AnkG) and MAP2 and imaged with a Zeiss LSM900 laser-scanning confocal microscope equipped with a 63x PlanApo NA 1.4 objective. Image stacks consisted of 6-8 optical sections acquired at 0.37 μm z-intervals. Z-stacks were projected into single images using maximum intensity projection in Fiji/ImageJ. The AIS was identified by AnkG immunoreactivity, and the neuronal soma was defined based on DAPI staining. A line region of interest (ROI) was then drawn from the soma along the AnkG-positive axon to delineate the AIS, following the method described by Grubb & Burrone (2010) [38]. Fluorescence intensity profiles along the ROI were smoothed over 1 μm intervals using SigmaPlot 12.5 and normalized to the maximal AnkG fluorescence, which was set to 100%. The proximal and distal boundaries of the AIS were defined as the points at which the normalized AnkG signal declined to 33% of the maximum at each end. AIS length was calculated as the distance between these two points, and the distance from the soma was defined as the distance between the soma and the proximal boundary.

### Calcium imaging

Hippocampal neurons were plated on poly-L-lysine–coated 25 mm round coverslips at a density of 20,000 cells/cm² and infected at DIV4 with lentiviral particles encoding mRuby2-jGCaMP8s, enabling expression of the calcium indicator jGCaMP8s [39] and the calcium-insensitive fluorescent protein mRuby2 [40], both under the control of CaMKIIα promoters. AAVs were added at DIV7. At DIV15, cultures were washed and incubated for 15 min at 37 °C in pre-warmed HBSS containing calcium and magnesium, and then transferred to Attofluor™ imaging chambers. Imaging was performed at 37 °C using a Zeiss Axiovert 200M epifluorescence microscope equipped with a 25× multi-immersion objective (NA 0.8), a CoolLED pE-4000 illumination system, and a PCO Edge 4.2 monochrome camera. jGCaMP8s fluorescence was recorded using 470/20 nm excitation and 525/50 nm emission filters. mRuby2 fluorescence was recorded using 550/15 nm excitation and 580/30 nm emission filters. Images were acquired at 10 Hz for 5 min. Overall fluorescence signals was background-subtracted using mean intensity values from cell-free regions, and individual neurons (soma plus proximal dendrites) were selected as regions of interest (ROIs). jGCaMP8s fluorescence was normalized to mRuby2 fluorescence to correct for motion artifacts and expression levels. Calcium signals were expressed as ΔF/F, where F represents the mean baseline fluorescence and ΔF the change in fluorescence at each time point relative to F. For pharmacological experiments, spontaneous activity was recorded for 1 min before drug application. Oscillation frequency and amplitude were quantified from ΔF/F traces obtained between 2 and 4 min after the start of image acquisition. Image processing and data analysis were performed using Fiji/ImageJ.

### Viability and survival assays

#### MTT assay

Primary hippocampal neuron cultures were seeded in 24-well plates and analyzed at DIV16. Thiazolyl Blue Tetrazolium Bromide (MTT; Sigma) was added to each well to a final concentration of 0.5 mg/ml. Cultures were incubated for 30 min at 37 °C to allow reduction of MTT to insoluble formazan crystals. Formazan was then solubilized in acidified isopropanol (isopropanol containing 0.04 N HCl), and absorbance was measured at 450 nm using a FluoStar Optima microplate reader (BMG Labtech). <u>LDH assay</u>: Cell membrane damage and cytotoxicity were assessed using a lactate dehydrogenase (LDH) release assay. At DIV16, 2 mM iodonitrotetrazolium chloride, 3.2 mM β-nicotinamide adenine dinucleotide (NAD, sodium salt), 160 mM lithium lactate and 15 μM 1-methoxyphenazine methosulfate (MPMS) in 0.2 M Tris–HCl (pH 8.2) were added to conditioned culture medium (NB or NB+) from hippocampal neurons. After incubating for 30 min at room temperature in the dark, the reaction was stopped by adding 1 M acetic acid, and LDH activity was quantified by measuring absorbance at 490 nm using a FluoStar Optima microplate reader (BMG Labtech). Data from LDH and MTT assays were exported to Excel for analysis and normalized to the corresponding control condition in each experiment.

#### Live-dead assay

Neuronal viability was assessed using a live–dead assay in primary hippocampal neuron cultures plated in 12-well plates. At DIV16, cultures were incubated in complete Hank’s balanced salt solution containing 1 µg/mL Hoechst 33342 (total nuclei), 1 µM Calcein-AM (viable cells), and 5 µM propidium iodide (PI; dead cells) for 30 min at 37 °C. Cells were then washed with complete Hank’s solution and imaged using a Zeiss AxioObserver7 fluorescence microscope. For each experimental condition, 20 random fields were acquired using a 20× objective, yielding quantitative analysis of approximately 800–1,000 cells per condition.

### Electrophysiology

Glass coverslips containing hippocampal neuron cultures (DIV15–17) were transferred to an open recording chamber and continuously perfused with artificial cerebrospinal fluid (aCSF) containing (in mM): 119 NaCl, 2.5 KCl, 2.5 CaCl, 1.2 MgCl, 26 NaHCO, 1 NaH PO, and 11 glucose, adjusted to pH 7.4 and 290 ± 5 mOsm. The aCSF was continuously bubbled with carbogen (95% O, 5% CO ), and temperature was maintained at 29°C. Neurons were previously cultured in NB+ medium and infected with Ng-EGFP AAVs. For recordings of AMPA receptor–mediated miniature excitatory postsynaptic currents (mEPSCs), aCSF was supplemented with 100 µM AP5 (NMDA receptor antagonist), 100 µM picrotoxin (GABA_A_ receptor antagonist), and 1 µM TTX to block action potentials. Patch electrodes (4–6 MΩ), pulled from borosilicate glass and containing Ag/AgCl wires, were filled with an internal solution composed of (in mM): 115 CsMeSO, 20 CsCl, 10 HEPES, 2.5 MgCl, 4 Na ATP, 0.4 Na-GTP, 10 sodium phosphocreatine, 0.6 EGTA, and 10 lidocaine N-ethyl bromide, adjusted to pH 7.25 and 290 ± 5 mOsm. Cells were voltage-clamped at −60 mV and recorded for 4–6 min using a gap-free protocol. Signals were acquired with a MultiClamp 700B amplifier, a Digidata 1550B A/D converter, and pCLAMP software (Molecular Devices). mEPSCs were detected and analyzed using Clampfit (Molecular Devices). For recordings of spontaneous activity and neuronal synchrony, patch electrodes were filled with an internal solution containing (in mM): 115 K -gluconate, 20 KCl, 10 HEPES, 2 MgCl, 4 Na ATP, and 0.3 Na GTP, adjusted to pH 7.2–7.3 and 290 ± 5 mOsm. Spontaneous activity was recorded simultaneously from pairs of neurons in current-clamp mode for 4–6 min using a gap-free protocol. Data were acquired using the same electrophysiological setup and analyzed in MATLAB. Spike synchrony was quantified following the method described by Quiroga et al. (2002), using a coincidence window of ±2.5 ms.

### Statistical analysis

Data were analyzed using GraphPad Prism 8. For parametric comparisons, Student’s t-test was used for two groups, and one-way or two-way ANOVA with Bonferroni post hoc correction for multiple groups. For non-parametric comparisons, the Mann–Whitney test or Kruskal–Wallis test was applied for two or more groups, respectively. Results are presented as mean ± SEM from at least three independent experiments. Statistical significance is defined as follows: * P < 0.05, ** P < 0.01, *** P < 0.001, **** P < 0.0001. The absence of asterisks indicates non-significant differences (ns).

## Results

### Neurogranin Regulates Neuronal Structure and Connectivity

Fig. 1a summarizes the effects of Ng expression and knockdown on Ng protein levels, assessed by Western blotting and by quantifying the proportion of Ng-positive neurons in DIV15 cultures. As shown, baseline Ng expression in cultured hippocampal neurons was low -approximately fivefold lower than in adult hippocampus-consistent with our previous observations [31]. Infection with AAV-Ng restored Ng abundance to adult levels and markedly increased the fraction of Ng-expressing neurons, from ∼18% to more than 60%. In the AAV-Ng construct, Ng expression is driven by the CaMKIIα promoter, ensuring selective expression in excitatory neurons. In contrast, shRNA-mediated knockdown reduced Ng protein to barely detectable levels, whereas the scrambled shRNA had no effect. These changes were consistently observed in both biochemical and cellular measurements. Thus, while endogenous Ng expression in cultured neurons remains far below adult levels, AAV-mediated expression effectively re-establishes physiologically relevant Ng abundance. This limited endogenous expression provides a useful baseline for evaluating Ng function, and together, these manipulations establish a robust framework for analyzing the consequences of Ng expression during neuronal development.

**Fig. 1.**
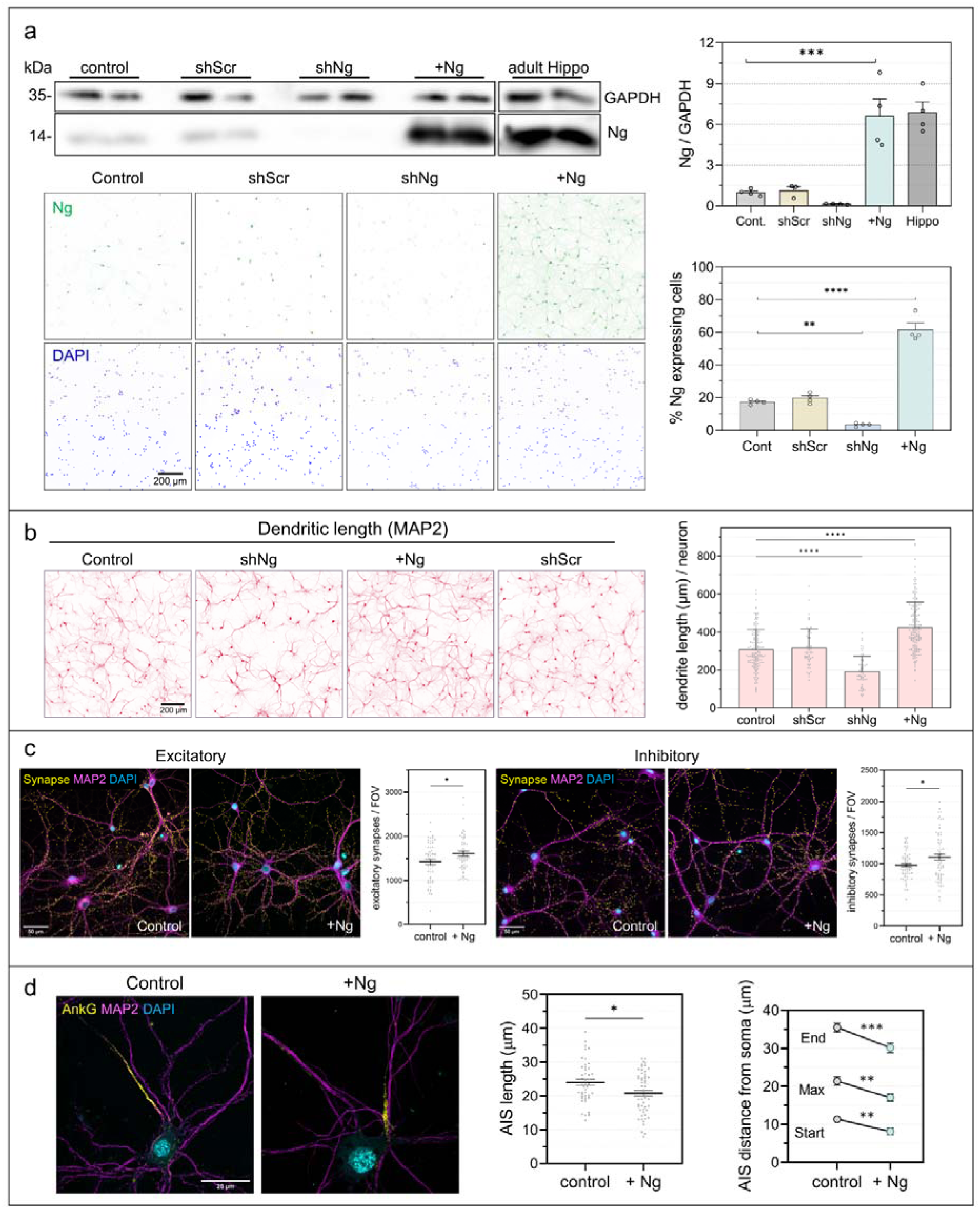
Ng expression modulates neuronal morphology and synapse density. Primary hippocampal neurons were infected with AAV-Ng at DIV7 or with AAV-shNg or scrambled shRNA (AAV-shScr) at DIV10 and analyzed at DIV15. (a) Ng protein levels were assessed by Western blot and normalized to GAPDH (upper panel; n = 4). The proportion of Ng-positive neurons was determined by fundingimmunofluorescence using DAPI to identify total cells (lower panel; n = 4). (b) Dendritic length per neuron, quantified for each field of view, increased following Ng expression. (c) Synaptic density was analyzed by co-localization of excitatory (vGluT1/PSD95) and inhibitory (GAD65/Gephyrin) pre- and postsynaptic markers. Data in histograms are total number of synapses per field of view (0.11 mm^2^). Ng expression increased the number of excitatory and inhibitory synaptic contacts (yellow puncta; n = 52–58 images). (d) The axon initial segment (AIS) was labeled with anti-ankyrin G (AnkG). Total AIS length (left graph; n = 25 neurons per condition) and distances from soma to AIS onset, peak intensity, and distal end (right graph) were quantified using DAPI to locate the soma. Widefield images (a–c) were acquired on a Zeiss Axiovert 200M microscope with 10× (A, B) or 40× (c) objectives; AIS images (d) were obtained using a Zeiss LSM900 confocal microscope with a 63× objective.

Morphological analysis revealed pronounced structural changes following both Ng expression and knockdown (Fig. 1b). AAV-Ng significantly enhanced dendritic growth, whereas shNg markedly reduced it. Quantification of total dendritic length per neuron confirmed these effects: dendritic length was substantially decreased by shNg, significantly increased by AAV-Ng, and unaffected by scrambled shRNA. We next examined synaptic number in control and AAV-Ng–infected cultures by measuring the colocalization of pre- and post-synaptic markers (Fig. 1c). Ng expression resulted in a clear increase in synapse number, both of excitatory and inhibitory synapses, indicating that Ng promotes synaptogenesis and synapse stabilization during neuronal development and maturation in vitro. Finally, we analyzed the axon initial segment (AIS) in control and AAV-Ng–infected neurons. As shown in Fig. 1d, Ng expression slightly shortened the AIS and shifted it to a position closer to the soma. This AIS remodeling most likely represents an adaptive response to the increased connectivity associated with Ng expression.

Together, these results show that restoring Ng to adult levels in excitatory neurons induces coordinated structural remodeling, including enhanced dendritic growth, increased synapse number, and adaptive repositioning of the AIS, whereas Ng depletion produces the opposite effects. These findings identify Ng as a critical determinant of neuronal connectivity during development and maturation and provide a foundation for subsequent functional analyses.

### Neurogranin Increases Spontaneous Activity and Network Synchronization

Next, we examined how Ng expression affects endogenous neuronal activity. Hippocampal neurons in culture begin firing action potentials from approximately DIV7 onwards [41, 42]. However, robust spontaneous activity is typically observed only at high plating densities, which are incompatible with the low-density conditions required for detailed morphological analyses. Using jGCaMP8, a genetically encoded calcium indicator, we first assessed the influence of plating density on spontaneous activity (Fig. 2a). At the density routinely used for our structural studies (20,000 cells/cm^2^), neurons exhibited no detectable activity in NB medium and only sparse, low-frequency events in NB+ medium, which promotes enhanced neuronal activity.

**Fig. 2.**
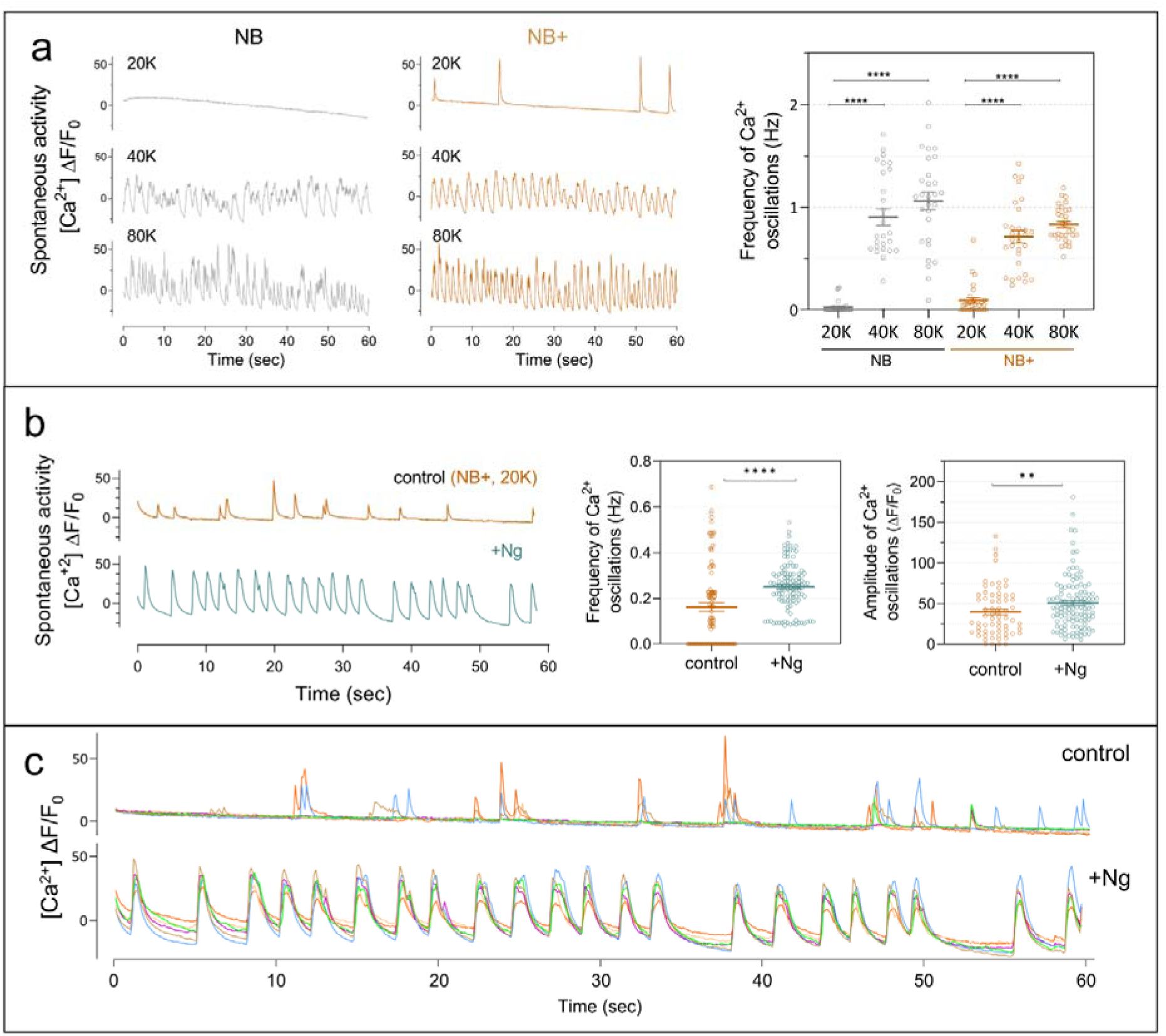
Ng expression enhances spontaneous calcium oscillations in hippocampal neurons. (a) Representative spontaneous Ca² oscillations detected using the jGCaMP8s biosensor in DIV15 hippocampal neurons cultured in NB or NB+ medium at different plating densities. Higher neuronal density increased the frequency of Ca² oscillations in both media. (b) Representative traces of spontaneous calcium activity in NB+ medium show increased frequency and amplitude of oscillations in Ng-expressing neurons (n = 100 neurons). (c) Synchronous Ca² activity across all neurons in the same field was observed following Ng expression.

In contrast, higher densities (40,000 and 80,000 cells/cm^2^) displayed robust spontaneous activity, consistent with previous observations [42]. Given that Ng expression enhances dendritic length and synaptic number, we focused on low-density cultures (20,000 cells/cm^2^) to assess its functional impact. Under these conditions, neurons expressing Ng exhibited markedly higher activity than controls, with increased frequency and amplitude of calcium transients (Fig. 2b). Moreover, Ng expression promoted strong synchronization of activity across neurons (Fig. 2c), consistent with enhanced synaptic connectivity. These results show that Ng expression is sufficient to robustly enhance endogenous neuronal activity and promote network synchronization in low-density cultures, indicating that Ng-driven structural changes translate into functional strengthening of synaptic coupling, potentially through altered network-level excitability.

To complement the calcium-imaging experiments and take advantage of higher temporal resolution, we performed whole-cell patch-clamp recordings. Electrophysiological recordings at DIV18 showed that neurons expressing Ng fired action potentials at a higher rate than control neurons (Fig. 3a). Detailed analysis showed that spikes frequently occurred in bursts or short high-frequency trains, with both burst incidence and the number of action potentials per burst significantly increased in Ng-expressing neurons (Fig. 3b). Paired recordings further revealed a marked enhancement of spike synchrony, as determined using a ±2.5-ms coincidence window (Fig. 3c). Despite this elevated firing and synchronization, Ng expression was associated with changes in intrinsic membrane properties indicative of reduced excitability. Ng-expressing neurons displayed a significantly more hyperpolarized resting membrane potential (Fig. 3d), and miniature excitatory postsynaptic currents (mEPSCs) exhibited lower frequency and amplitude compared with controls (Fig. 3e). Together, whole-cell recordings reveal increased firing, burst activity, and spike synchrony despite a more hyperpolarized resting membrane potential and reduced mEPSC amplitude, arguing against a simple increase in intrinsic neuronal excitability. Instead, by promoting dendritic growth and increasing synaptic number, Ng likely facilitates a reorganization of network-level signaling that supports efficient and coordinated activity through strengthened synaptic connectivity, even at low neuronal densities.

**Fig. 3.**
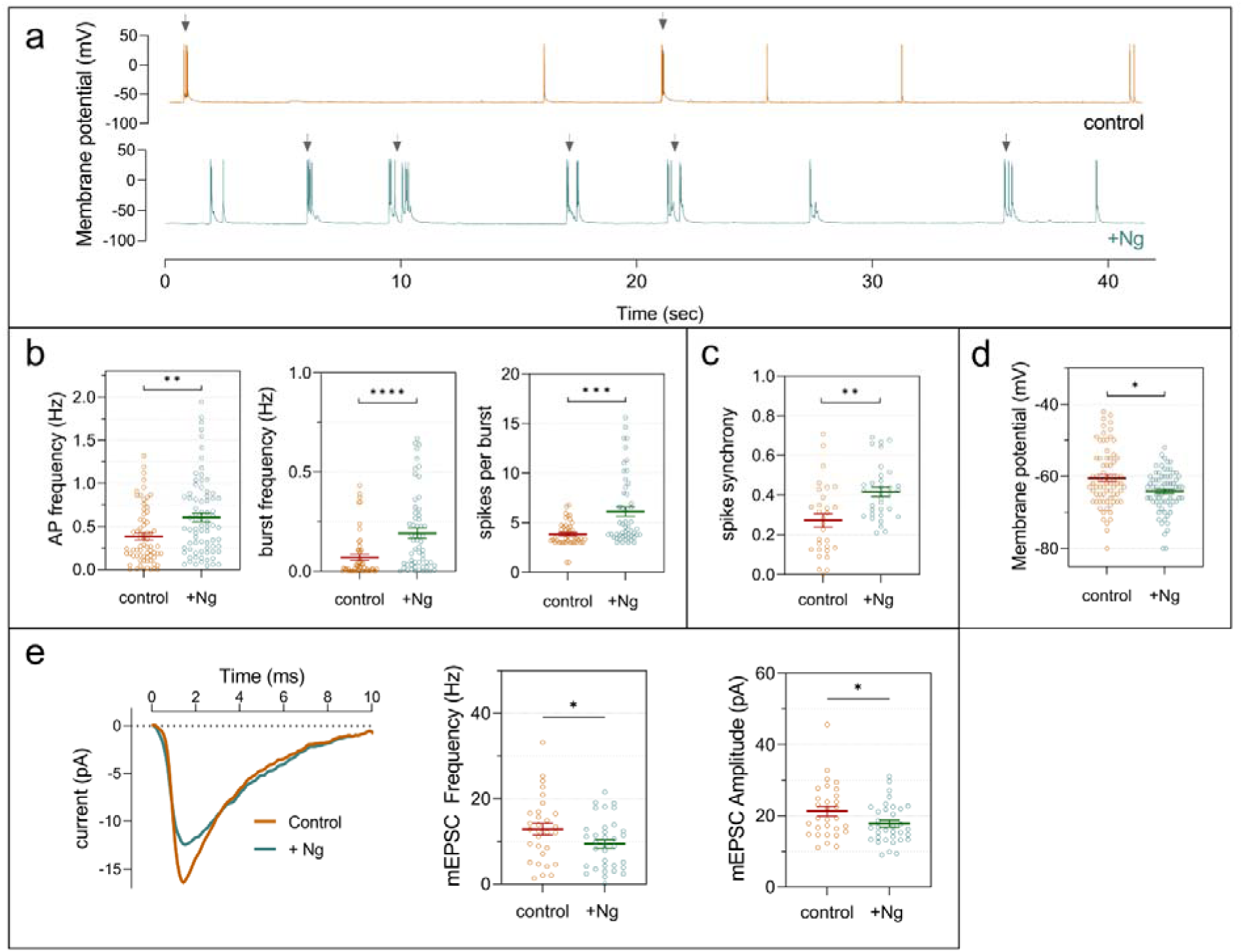
Ng expression increases spontaneous electrical activity and network synchronization. Hippocampal neurons cultured in NB+ medium were analyzed at DIV18 by whole-cell patch-clamp. (a) Representative 40-s current-clamp traces showing spontaneous action potential (AP) bursts (arrows) in control and Ng-expressing neurons. (b) Quantification of AP frequency, burst frequency, and APs per burst (n = 70– 75 neurons). (c) Spike synchrony was assessed by simultaneous recordings from neuron pairs as described in Materials and Methods. (d) Resting membrane potential was more hyperpolarized in Ng-expressing neurons. (e) Miniature excitatory postsynaptic currents (mEPSCs) were recorded in the presence of 1 µM TTX and 10 µM bicuculline. Representative averaged mEPSCs are shown (left), and histograms depict frequency and amplitude distributions (right).

### Cellular and Synaptic Mechanisms of Neurogranin Action

To elucidate the mechanisms underlying Ng-dependent enhancement of network activity, we monitored intracellular Ca^2+^ dynamics in response to pharmacological manipulations and analyzed the expression of key synaptic and signaling proteins. First, we confirmed that Ng-induced increases in spontaneous activity require action potential firing: application of tetrodotoxin (TTX), a voltage-gated sodium channel blocker, completely abolished the elevated Ca^2+^ transients, indicating dependence on neuronal firing and synaptic activity (Fig. 4a). We then examined the role of ionotropic glutamatergic receptors. Short-term NMDA receptor blockade with AP5 selectively reduced the amplitude of high-calcium events without affecting firing frequency (Fig. 4b), whereas AMPA receptor inhibition with NBQX decreased both event amplitude and firing frequency (Fig. 4c). These results indicate that NMDARs and AMPARs contribute to the magnitude and frequency of calcium signals. Notably, mGluR5 inhibition with MPEP strongly reduced both frequency and amplitude of calcium transients (Fig. 4d). Given that mGluR5 activates Gq/11-coupled phospholipase C signaling, leading to IP_3_-mediated Ca^2+^ release from intracellular stores, we tested the IP3 receptor inhibitor 2-APB, which markedly diminished calcium event frequency and amplitude (Fig. 4e), confirming the involvement of intracellular Ca² signaling in high-amplitude events. These findings indicate that Ng enhances network activity through coordinated mechanisms involving ionotropic glutamatergic signaling, and mGluR5/IP3-mediated intracellular calcium mobilization, providing a mechanistic basis by which Ng strengthens synaptic connectivity and network dynamics.

**Fig. 4.**
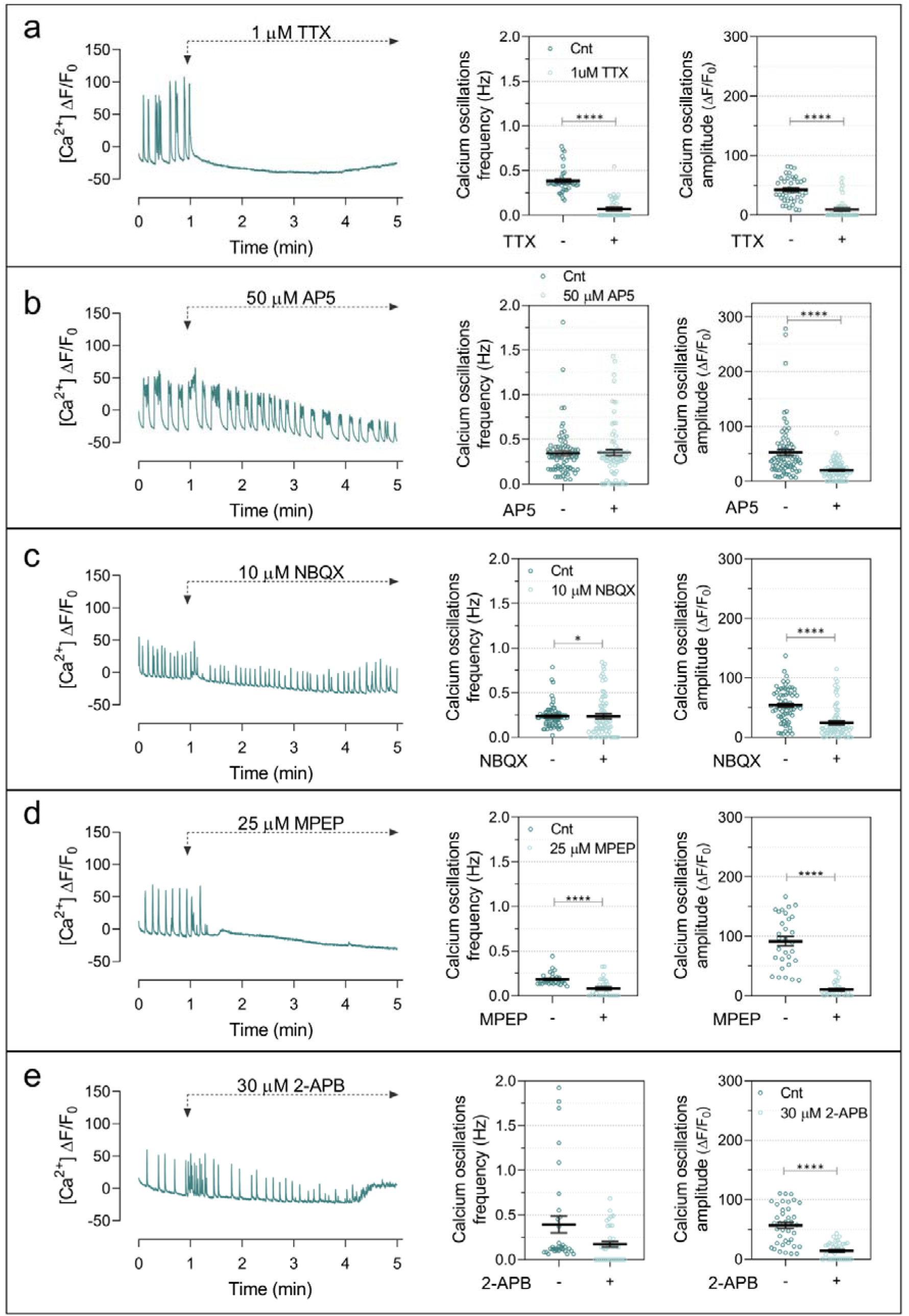
Synaptic signaling mediates Ng-induced increases in spontaneous calcium activity. Spontaneous DIV15 hippocampal neurons infected with AAV-Ng and maintained in NB+ medium from DIV7 were monitored for spontaneous Ca² oscillations. Baseline activity was recorded for 1 min, followed by acute application of pharmacological agents: 1 µM TTX (a), 50 µM AP5 (b), 10 µM NBQX (c), 25 µM MPEP (d), or 30 µM 2-APB (e). Ca² activity was then recorded for 2 min, starting 1 min after treatment. Representative calcium traces are shown, and histograms compare oscillation frequency and amplitude before and after drug application. Between 30 and 90 cells were analyzed per condition.

Although Ng enhances spontaneous activity through glutamatergic signaling, the underlying molecular mechanisms remained unclear. To address this, we examined the impact of Ng expression on key components of Ca^2+^/CaM-dependent signaling and synaptic function. Ng binds CaM and is proposed to act as a CaM-sequestering protein [29, 43]. In both NB and NB+ media, Ng expression significantly increased total CaM levels (Fig. 5a), with a more pronounced effect in NB medium, which exhibits lower baseline activity and reduced neuronal survival. Analysis of Ng mutants showed that only Ng-wt and the CaM-binding-competent mutant Ng-Ser36Ala increased CaM levels, whereas CaM-binding-deficient mutants (Ser36Glu and Ile33Gln) had no effect, supporting the notion that Ng sequesters CaM and that compensatory mechanisms may adjust CaM expression or turnover in response to changes in free (apo-)CaM. We next examined Ca^2+^/CaM-dependent protein kinase II (CaMKII), a central mediator of synaptic plasticity and LTP [44]. Ng expression reduced total CaMKII levels in both NB and NB+ media (Fig. 5b), yet substantially increased the phospho-Thr286 CaMKII to total CaMKII ratio (pCaMKII/CaMKII) (Fig. 5c), indicating a shift toward a higher relative activation state. Analysis of Ng mutants showed that only CaM-binding-competent forms of Ng were able to elevate this ratio (Suppl Fig. 1). In contrast, the levels of the phosphatase Calcineurin (A subunit, CaN-A) remained unchanged under all conditions tested (Fig. 5b, Suppl. Fig. 2). These findings are consistent with enhanced network activity in Ng-expressing neurons and suggest more efficient CaM-dependent signaling despite reduced CaMKII abundance. We then assessed ionotropic glutamate receptors. Ng expression selectively reduced total and surface levels of NMDA receptor subunits GluN1 and GluN2B, while GluN2A remained unchanged (Fig. 5d,g). Similarly, total and surface levels of the AMPA receptor subunit GluA1 were decreased, whereas GluA2 and mGluR5 levels were unaffected (Fig. 5e-g). Surface expression closely paralleled total protein levels, indicating that Ng selectively downscales excitatory receptor abundance. Together, these observations indicate a coordinated reorganization of Ca² /CaM-dependent signaling and synaptic composition. By sequestering CaM, Ng shifts the balance between free and bound CaM, enhancing CaMKII activation while simultaneously inducing a homeostatic downscaling of specific ionotropic glutamate receptors. This adaptive response maintains efficient network signaling and synchronization while restraining postsynaptic excitability, and balances increased connectivity with controlled excitatory drive.

**Fig. 5.**
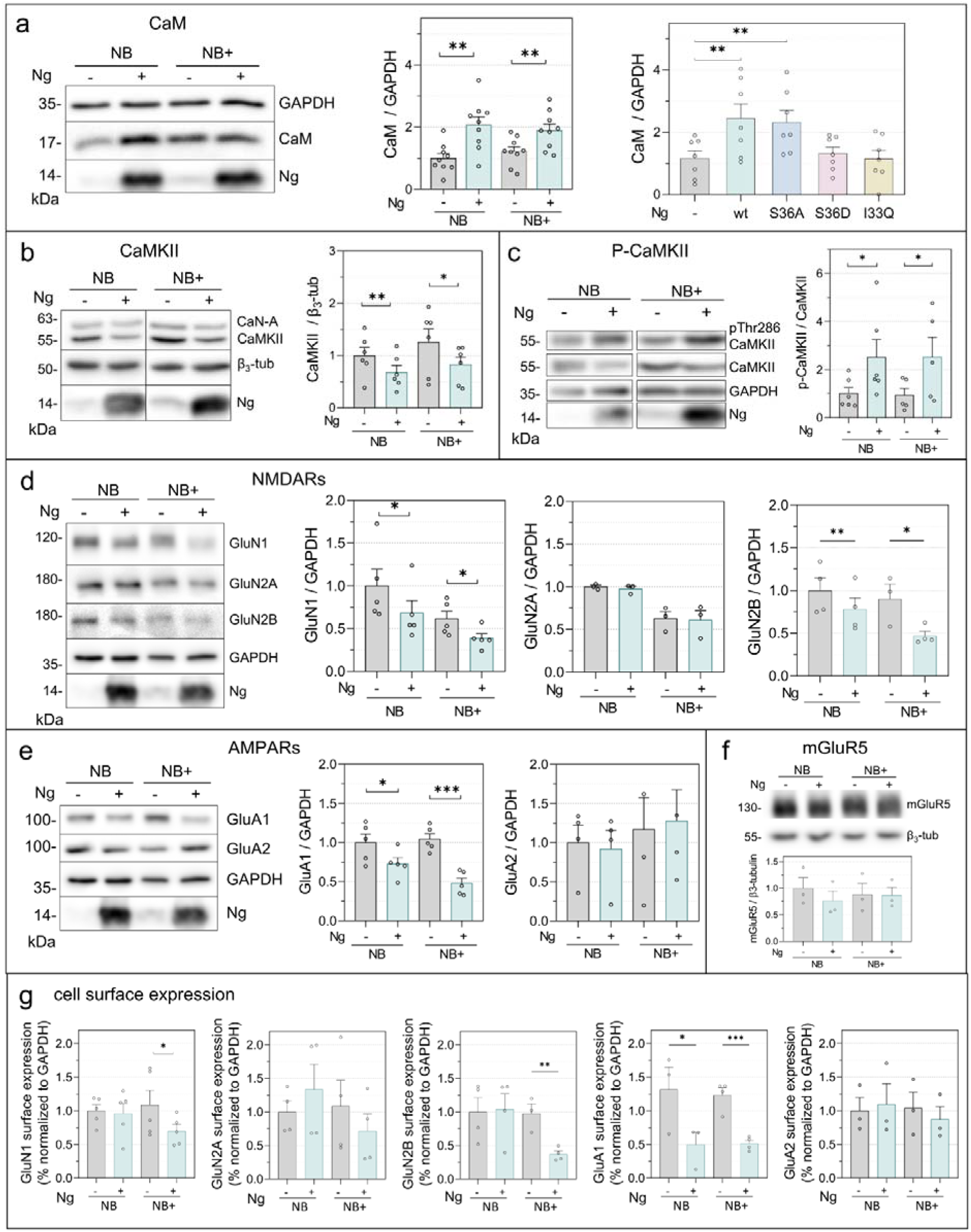
Ng expression alters synaptic protein abundance and surface expression. Primary hippocampal neurons were infected with AAV-Ng and maintained in NB or NB+ medium from DIV7. Protein extracts at DIV16 were analyzed by Western blot, and levels were normalized to GAPDH or β3-tubulin. Histograms show mean normalized values relative to non-infected neurons in NB. (a–c) CaM, total CaMKII, and phospho-CaMKII (Thr286) were analyzed. (d–e) NMDA receptor subunits (GluN1, GluN2A, GluN2B) and AMPA receptor subunits (GluA1, GluA2) were assessed. (f) mGluR5 levels were analyzed. (g) Surface expression of NMDA and AMPA receptors was determined by sulfo-NHS-biotin labeling followed by streptavidin–agarose isolation. GAPDH served as a control for the input fraction (n = 4).

### Neurogranin Expression enhances Neuronal Viability and Survival

Given the pronounced effects of Ng on neuronal activity and network synchronization, we next examined whether these changes are detrimental or beneficial for neuronal viability. Metabolic activity was first assessed using the MTT assay (Fig. 6a). In NB medium, Ng expression significantly increased mitochondrial reducing activity, whereas Ng knockdown caused only a modest, non-significant reduction. In NB+ medium, which supports higher baseline metabolic activity, the most prominent effect was observed upon Ng knockdown, which led to a clear reduction in metabolic activity. We then evaluated membrane integrity and cytotoxicity using the lactate dehydrogenase (LDH) release assay (Fig. 6b). Ng expression consistently reduced LDH release, while Ng knockdown increased LDH levels in both NB and NB+ media, indicating that Ng confers protection against membrane damage and cell death. Neuronal viability was further examined using the calcein-AM/propidium iodide (PI) assay (Fig. 6c). Quantification of live (calcein-positive) and dead (PI-positive) cells revealed that Ng expression increased the proportion of viable neurons and reduced necrotic cell death in both culture conditions. Notably, this protective effect was even more pronounced at DIV23, a time point associated with a substantial decline in neuronal viability (Suppl. Fig. 3). Together, these results demonstrate that, despite markedly enhancing neuronal activity and synchronization, Ng robustly promotes neuronal viability and survival. Importantly, the levels of Ng achieved by AAV-mediated expression closely match those found in adult hippocampal neurons in vivo, underscoring the physiological relevance of these effects.

**Fig. 6.**
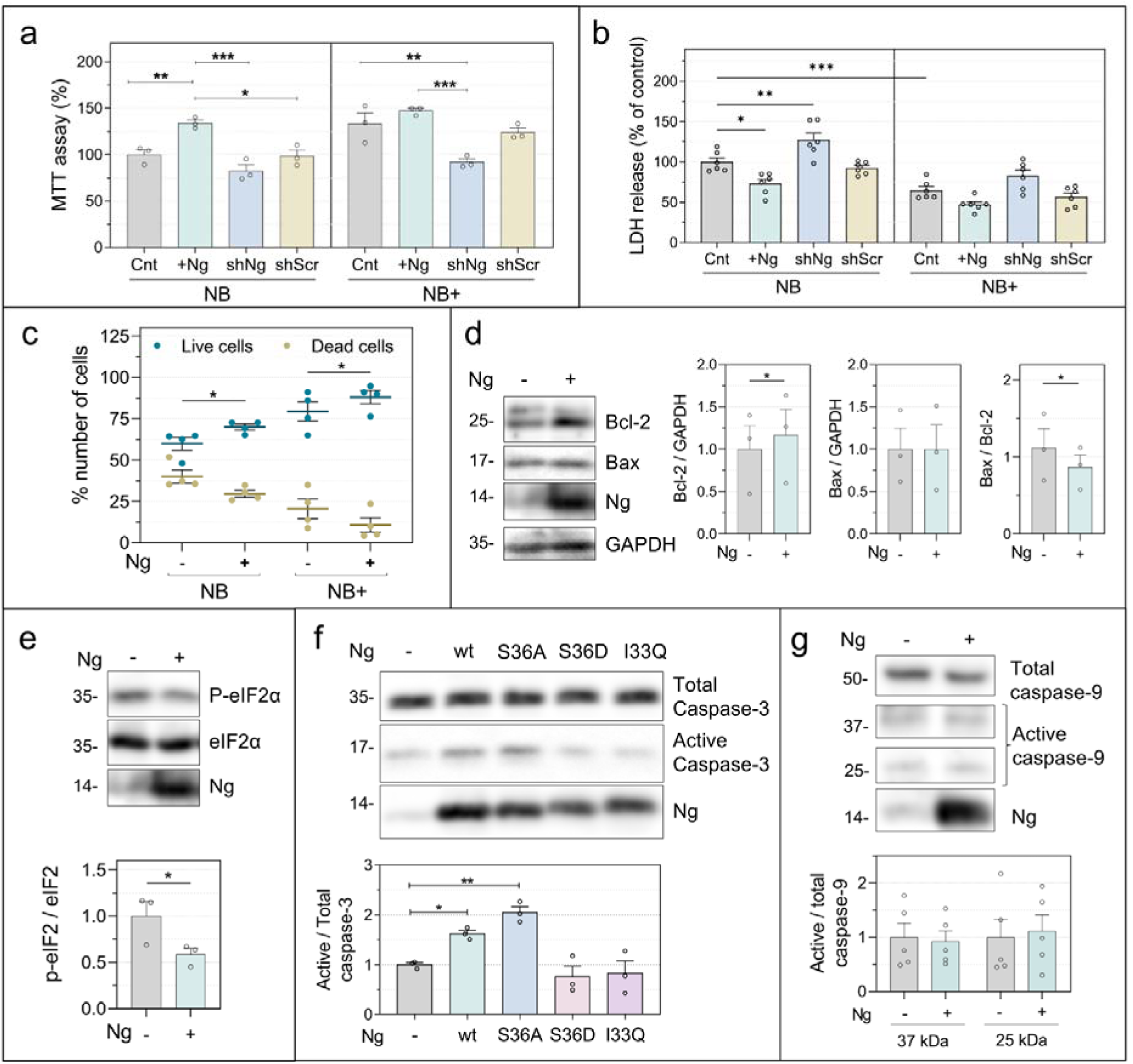
Ng expression promotes neuronal viability and stress resilience. Primary hippocampal neurons were infected with AAV-Ng or AAV-shNg and maintained in NB or NB+ medium from DIV7. (a) Mitochondrial metabolic activity was assessed at DIV16 using the MTT assay. (b) Cytotoxicity and membrane integrity were assessed using the lactate dehydrogenase (LDH) release assay. (c) Neuronal viability was evaluated using a combined Calcein-AM / Propidium Iodide (PI) / Hoechst 33342 assay at DIV16. Live cells were identified by Calcein-AM fluorescence, dead cells by PI staining, and total cell number by Hoechst 33342 labeling (1 µg/mL). (d) Expression of the anti-apoptotic protein Bcl-2 and the pro-apoptotic protein Bax was analyzed in neuronal extracts at DIV16 and normalized to GAPDH. (e) Phosphorylation of eIF2α at Ser51, an indicator of integrated stress response activation, was analyzed by Western blot and normalized to total eIF2α levels. (f–g) Levels of active (cleaved) caspase-3 and caspase-9, respectively, were quantified and normalized to their respective total protein levels. Data are presented as mean values ± SEM relative to control conditions (n = 4 independent experiments).

To investigate the mechanisms underlying the enhanced neuronal viability, we next examined markers of cellular stress and key regulators of the intrinsic apoptotic pathway. Ng expression did not alter the levels of the pro-apoptotic protein Bax, which promotes mitochondrial outer membrane permeabilization, but produced a modest yet significant increase in the anti-apoptotic protein Bcl-2, resulting in a reduced Bax/Bcl-2 ratio (Fig. 6d). In parallel, Ng expression significantly decreased phosphorylation of eIF2α at Ser51, a central effector of the integrated stress response, consistent with attenuated cellular stress signaling and improved adaptive capacity (Fig. 6e). We then assessed caspase-3 activation, a hallmark effector of apoptotic pathways but also a mediator of non-apoptotic neuronal remodeling [45]. Neurons expressing CaM-binding Ng variants (Ng-wt and Ng-Ser36Ala) exhibited a moderate increase in active caspase-3 levels (Fig. 6f). Notably, this increase was not associated with reduced neuronal viability, arguing against apoptotic execution. Instead, controlled caspase-3 activation has been linked to synaptic plasticity and structural remodeling in neurons [46–51]. The restriction of this effect to CaM-binding Ng variants, which also elevate total CaM levels, supports the involvement of Ca² /CaM-dependent signaling in this response. Consistent with a non-apoptotic mechanism, Ng expression did not affect the levels of active caspase-9, an initiator of the intrinsic apoptotic pathway (Fig. 6g). Collectively, these findings indicate that Ng enhances neuronal survival by reducing cellular stress and engaging regulated, activity-dependent remodeling pathways, rather than triggering apoptotic cell death.

## Discussion

In this study, we identify Ng as a central regulator of neuronal activity, network synchronization, and long-term neuronal viability through its capacity to modulate Ca^2+^/CaM signaling. By integrating immunofluorescence, functional calcium imaging, electrophysiology, quantitative analysis of synaptic markers, and multiple viability assays, we show that Ng expression drives morphological maturation, enhances spontaneous neuronal activity and network synchrony, and concomitantly promotes metabolic competence, stress resilience, and neuronal survival. Importantly, these effects depend on Ng’s ability to bind CaM, establishing a direct mechanistic link between activity-dependent Ca^2+^ signaling and coordinated structural, functional, and homeostatic adaptations at the cellular and network levels.

Using cultured hippocampal neurons as a model system, we examined the impact of Ng expression at three levels: neuronal morphology, network activity, and neuronal viability. Notably, AAV-mediated Ng expression achieved protein levels comparable to those observed in the adult hippocampal tissue. Moreover, expression was driven by the CaMKIIα promoter, thereby restricting Ng expression spatially and temporally to excitatory neurons from DIV7 onward. This experimental design enabled the assessment of Ng function within a physiologically relevant expression range and a defined developmental window.

Dendritic arbor architecture is a key determinant of neuronal information processing, as neurons dynamically adapt their structure to their functional roles within neural circuits [52]. Ng expression promoted increased dendritic complexity and synaptic number, yielding morphologies that more closely resemble those of mature hippocampal neurons in vivo. In vitro, enhanced neuronal morphology and connectivity are typically associated with favorable trophic conditions and robust metabolic support, and are regulated by neuronal activity, intracellular Ca² signaling, and synaptic input [53, 54]. In hippocampal neuron cultures, synaptogenesis begins toward the end of the first week in vitro, increases during the second week, and peaks during the third week [55]. As shown here and in previous studies [56], both spontaneous activity and dendritic development are strongly influenced by plating density, which determines network connectivity. Under the relatively low-density conditions used in this study (20,000 cells/cm²), where dendritic growth and connectivity are otherwise limited, Ng expression markedly enhanced dendritic arborization and synaptic number. The accompanying increase in spontaneous activity is therefore likely to be a major driver of these structural changes. Collectively, these findings indicate that Ng facilitates developmental maturation programs that promote the emergence of integrated and functional neuronal networks. Consistent with this interpretation, it has been shown that Ng maintains glutamatergic balance by coordinating experience-dependent synapse elimination and promoting the conversion of silent synapses into functional, AMPAR-containing synapses, thereby optimizing circuit performance [57].

A substantial body of experimental and computational work indicates that Ng binds free CaM (apo-CaM) within dendritic spines and that this interaction -dynamically regulated by phosphorylation and oxidation-shapes local Ca² /CaM signaling during synaptic activity [22, 23, 58, 59]. By concentrating CaM at synapses, Ng is proposed to amplify activity-dependent Ca² signals such that each synaptic Ca² influx generates higher local Ca² /CaM levels, thereby facilitating the activation of downstream effectors with relatively low Ca² /CaM affinity, most notably CaMKII [60]. Sustained engagement of this mechanism is predicted to promote synaptic potentiation, synaptogenesis, and the expansion and stabilization of dendritic arbors. In parallel, Ng expression shifted the AIS closer to the soma, a structural change expected to lower spike threshold [38] and thus contribute to the enhanced firing activity observed.

Our findings are consistent with this framework of enhanced excitatory drive but significantly extend it. While Ng increased the relative activation state of CaMKII, as reflected by elevated Thr286 phosphorylation, it concurrently reduced total CaMKII abundance, downscaled specific ionotropic glutamate receptor subunits, hyperpolarized the resting membrane potential, and reduced mEPSC amplitude. Thus, Ng does not simply potentiate glutamatergic transmission; in addition, it induces a coordinated reorganization of CaM-dependent signaling and synaptic composition that supports elevated neuronal activity and efficient network synchronization while preserving overall excitatory balance. This pattern is characteristic of homeostatic plasticity [61], in which sustained increases in network activity elicit compensatory mechanisms that constrain synaptic strength and stabilize neuronal output. Notably, the increased phosphorylation of CaMKII observed in Ng-expressing neurons aligns with *in vivo* findings in Ng knockout mice, in which basal CaMKII autophosphorylation is reduced by approximately 50% [20]. Together, these complementary gain- and loss-of-function observations support a model in which Ng tunes the availability of Ca² /CaM to set the basal activation state of CaMKII.

Using calcium imaging to dissect the mechanisms underlying the Ng-dependent enhancement of neuronal activity, we found that Ca^2+^ transients strictly depend on action potential firing and are strongly attenuated by inhibition of ionotropic glutamate receptors, consistent with a synaptically driven process. Unexpectedly, inhibition of metabotropic glutamate receptors -specifically mGluR5-produced a profound suppression of Ca^2+^ activity, markedly decreasing both event frequency and amplitude. These findings indicate that spontaneous network activity relies not only on fast ionotropic transmission but also on metabotropic signaling and Ca² release from intracellular stores. Notably, Ng expression did not alter total mGluR5 protein levels, suggesting that enhanced receptor engagement, signaling efficiency, or downstream coupling - rather than increased receptor abundance-accounts for this effect. The increased prevalence of burst firing in Ng-expressing neurons is likely to favor mGluR5 activation, as these perisynaptic receptors require sustained or high local glutamate concentrations in the synaptic cleft for effective signaling. Together, these observations support a model in which Ng amplifies spontaneous network activity by promoting cooperative ionotropic–metabotropic signaling and intracellular Ca² mobilization, rather than by simply increasing excitatory synaptic drive.

Despite markedly altering neuronal firing patterns, Ng does not compromise neuronal viability or survival. Instead, it engages adaptive, non-apoptotic programs, as evidenced by increased metabolic activity (MTT assay), preserved plasma membrane integrity (LDH release), enhanced cell viability (Calcein/PI assay), reduced Bax/Bcl-2 ratios, and decreased phosphorylation of eIF2α at Ser51. Concurrently, Ng expression induces a moderate but significant increase in active caspase-3 that occurs independently of caspase-9 activation, indicating that the intrinsic mitochondrial apoptotic pathway is not engaged. This pattern aligns with well-documented non-apoptotic roles of caspase-3 in synaptic remodeling, dendritic restructuring, and neuronal differentiation [62–65, 50], rather than neurotoxicity. To date, no direct interaction between CaM and caspase-3 has been reported, nor did we observe evidence for direct coupling between Ca² /CaM signaling and caspase-3 activation in this context. On the contrary, CaM inhibition has been shown to promote caspase-3 activation [66], potentially via effects on intracellular Ca² homeostasis. This supports the idea that Ng-mediated CaM sequestration may indirectly favor localized, controlled caspase-3 activity. Previous studies have shown that NMDA receptor–dependent synaptic plasticity, including long-term depression and AMPAR internalization, require caspase-3 activity [67] and involve transient caspase-3 activation without progression to apoptosis. In this framework, the moderate caspase-3 activation observed in Ng-expressing neurons may contribute to synaptic downscaling, synapse pruning, and dendritic branch remodeling, coordinating structural plasticity with elevated network activity. Collectively, these findings support a model in which Ng functions not only as a synaptic CaM buffer but also as a molecular integrator that couples enhanced neuronal activity and connectivity to adaptive structural and survival responses. In this context, caspase-3 serves a sculpting role -refining synaptic and dendritic architecture-rather than a cytotoxic one [65], promoting network maturation and stabilization while preserving neuronal integrity.

Regarding Ng function during neuronal development, previous work from our group demonstrated that Ng expression promotes synapse formation in cultured neurons [31], and Han et al. [57] showed that Ng coordinates both the conversion of AMPAR-silent synapses into functional synapses and experience-dependent synapse elimination during critical developmental windows, thereby optimizing synaptic transmission and network balance. Consistent with these roles, Ng expression emerges relatively late in brain development [2], coinciding with periods of intense synaptogenesis and circuit remodeling rather than earlier stages of cell proliferation or migration. Notably, brain Ng levels exhibit a bimodal relationship with cognitive performance: reduced Ng is linked to cognitive deficits in several neurological and psychiatric conditions, including hypothyroidism [18, 68], schizophrenia [69], and Alzheimer’s disease [16] whereas elevated Ng levels in cerebrospinal fluid -reflecting synaptic loss and dysfunction-correlate with disease progression and early cognitive decline, particularly in Alzheimer’s disease [10, 13]. Collectively, these findings highlight Ng as a key regulator of synaptic development, circuit refinement, and the maintenance of cognitive function.

In summary, our findings indicate that Ng expression in cultured hippocampal neurons during the period of peak synaptogenesis and circuit maturation enhances morphological differentiation, increases synaptic connectivity, elevates neuronal activity, and strengthens network synchronization. These changes are accompanied by a compensatory downscaling of neuronal excitability and improved neuronal viability, suggesting that Ng drives maturation toward a state that more closely resembles in vivo physiological conditions. We propose that Ng supports the construction and stabilization of complex neuronal circuits with greater information-processing capacity while preserving excitatory balance and neuronal integrity. Although this study focuses on Ng function during neuronal development and maturation, and these conclusions cannot be directly extrapolated to mature adult neurons, Ng levels are known to decline progressively with aging [14], coinciding with cognitive deterioration. Furthermore, Ng loss in brain tissue, together with its concomitant increase in cerebrospinal fluid [70], is accelerated during the early stages of neurological disorders associated with cognitive impairment. Taken together with our findings, these observations suggest that strategies aimed at preserving or restoring Ng expression in excitatory neurons could help delay or mitigate cognitive decline.

## Acknowledgements

We thank the Advanced Light Microscopy Core Facility (SMOA) of the “Centro de Biología Molecular” (CBM) for assistance with the imaging studies. We thank “Fundación Ramón Areces” for providing institutional support to Centro de Biología Molecular (CSIC-UAM).

## Abbreviations

2-APB: (2-aminoethoxydiphenyl borate),
AAV: (adeno-associated virus),
AD: (Alzheimer’s disease),
AIS: (axon initial segment),
AnkG: (Ankyrin G),
AraC: (1-beta-arabino-furanosylcytosine),
BSA: (bovine serum albumin),
Ca^2+^: (calcium ion),
CaM: (calmodulin),
CaMKII: (calcium/calmodulin-dependent protein kinase II),
CSF: (cerebrospinal fluid),
DIV: (days in vitro),
DMEM: (Dulbecco’s modified Eagle’s medium),
E19: (embryonic day 19),
ECL: (enhanced chemiluminescence),
FBS: (fetal bovine serum),
HBSS: (Hank’s balanced salt solution),
LDH: (lactate dehydrogenase),
LTD: (long-term depression),
LTP: (long-term potentiation),
LVs: (lentiviral particles),
MCI: (mild cognitive impairment),
mEPSC: (miniature excitatory postsynaptic current),
MPEP: (2-methyl-6-(phenylethynyl)pyridine),
MPMS: (1-methoxyphenazine methosulfate),
MTT: (thiazolyl blue tetrazolium bromide),
NA: (numerical aperture),
NAD: (β-nicotinamide adenine dinucleotide),
NB: (neurobasal culture medium),
Ng: (Neurogranin),
NMDA: (N-methyl-D-Aspartate),
PA: (phosphatidic acid),
PFA: (paraformaldehyde),
PI: (propidium iodide),
PKC: (protein kinase C),
PLL: (poly-L-lysine),
ROI: (region of interest),
RT: (room temperature),
TTX: (tetrodotoxin)

## Statements and Declarations

### Authors Contributions

Elena Martínez-Blanco (EM-B) and Raquel de Andrés (RdA) performed material preparation, data acquisition, and analysis. Esperanza López-Merino (EL-M) and Elena Martínez-Blanco (EM-B) conducted the electrophysiology experiments under the supervision of Jose A. Esteban (JAE). F. Javier Díez-Guerra (F.J.D-G.) conceived and designed the study, contributed to data analysis, and wrote the manuscript. All authors commented on previous versions of the manuscript. All authors read and approved the final manuscript.

### Data Availability

The datasets generated and analyzed during the current study are available from the corresponding author on reasonable request

### Ethics approval

All procedures conducted during the study strictly adhered to the Spanish Royal Decree 1201/2005, which governs the protection of animals used in scientific research, as well as the European Union Directive 2010/63/EU concerning the welfare of animals in scientific contexts. Experimental protocols were approved by the Animal Experimentation Ethics Committee of the Centro de Biología Molecular (CEEA-CBMSO-23/247), the Research Ethics Committee of the Universidad Autónoma de Madrid (CEI-126-2594) and the Directorate-General for Agriculture, Livestock and Food of the “Comunidad de Madrid” (PROEX 106/19), ensuring that the highest standards of animal care and ethical compliance were maintained throughout the research.

---

**Supplementary Fig. 1.**
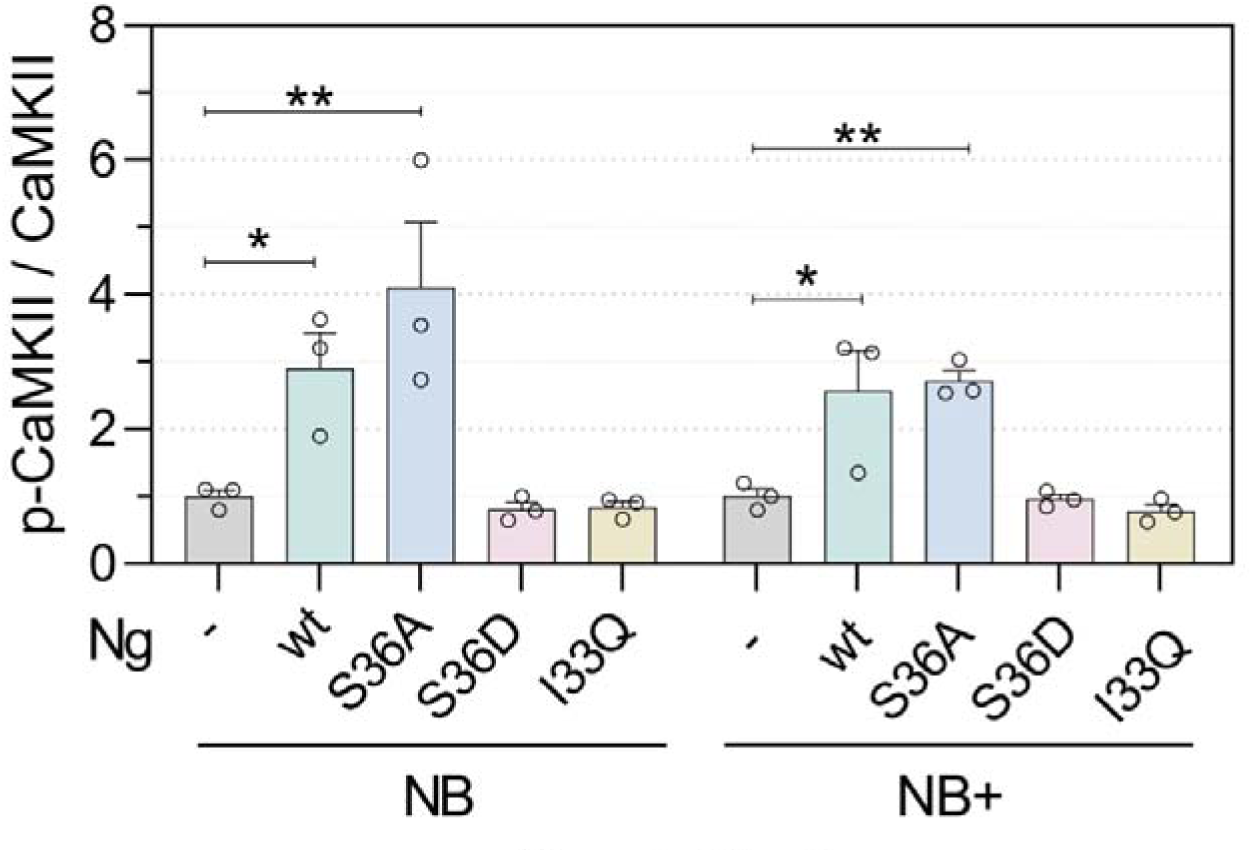
Effect of Ng mutants on the p-CaMKII/CaMKII ratio. Primary hippocampal neurons were infected at DIV7 with AAV-Ng wt or the indicated mutants and maintained in NB or NB+ media. Protein extracts were collected at DIV16 and analyzed by Western blot to quantify phosphorylated CaMKII and total CaMKII. The p-CaMKII/CaMKII ratios were calculated and normalized to non-infected neurons maintained in NB medium. Neurons expressing wild-type Ng or the S36A mutant showed a marked increase in the p-CaMKII/CaMKII ratio, whereas mutants deficient in calmodulin binding did not. (n = 3, mean ± SEM).

**Supplementary Fig. 2.**
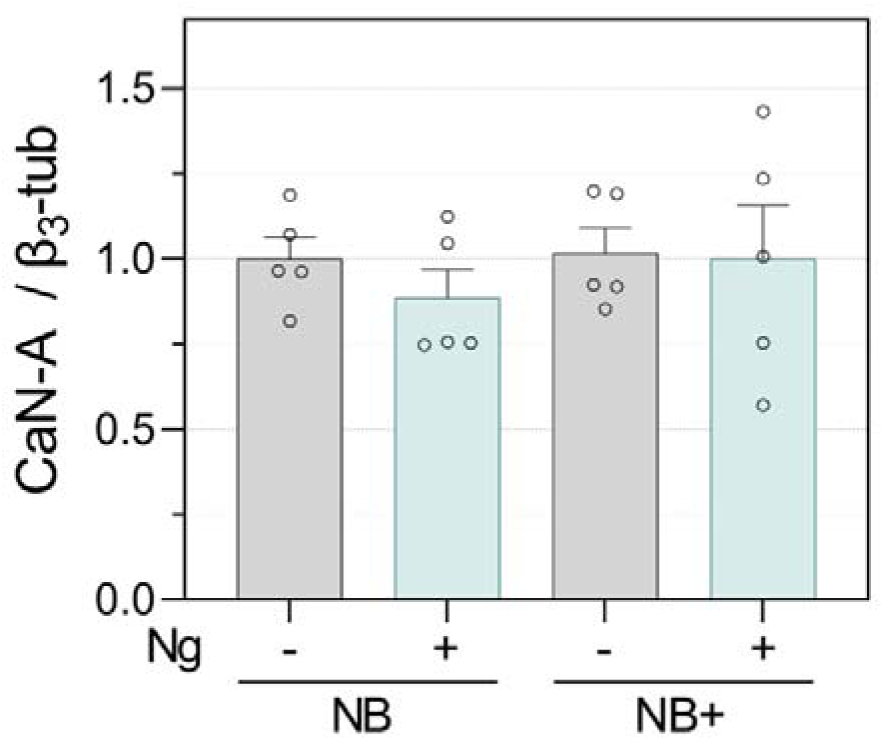
Effect of Ng on Calcineurin-A levels. Cultured hippocampal neurons infected with AAV-Ng at DIV7 and maintained in NB or NB+ medium were extracted at DIV16 and analyzed by Western blot. CaN-A levels were normalized to β3-tubulin. Histogram show mean normalized values relative to non-infected neurons in NB. (n = 5, mean ± SEM).

**Supplementary Fig. 3.**
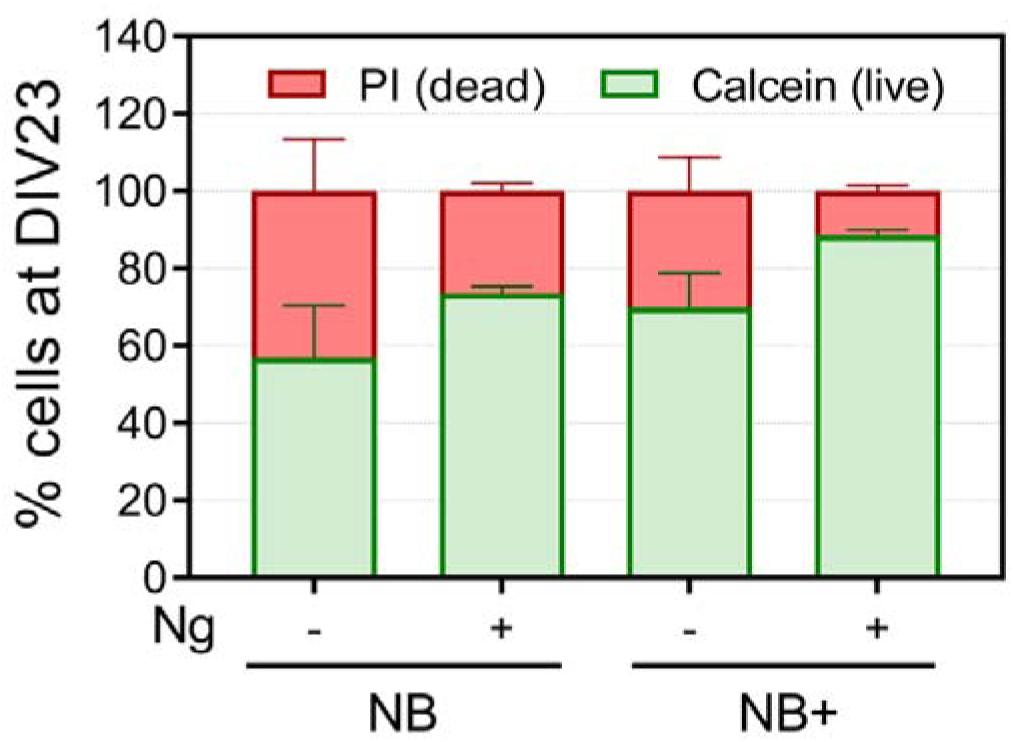
Effect of Ng expression on cell viability in DIV23 cultures. Cell viability was assessed at DIV23 using Calcein-AM, propidium iodide, and Hoechst 33342, as described in Fig. 6c. In Ng-expressing cultures exhibited a higher number of live cells compared with matched control cultures. (n = 3, mean ± SEM).

**Supplementary Table 1.** Plasmids developed and used in this study.

| Plasmids | Backbone | Insert | promoter | expression | tag |
| --- | --- | --- | --- | --- | --- |
| **pLV-mRuby2-jGCaMP8s** | pLOX | mRuby & jGCaMP8s | CaMKIIα0.4 | Lentiviral | mRuby2 |
| **pAAV-Ng** | **pAAV | Ng | CaMKIIα0.4 | AAV2 | |
| **pAAV-Ng-S36A** | pAAV | Ng-S36A | CaMKIIα0.4 | AAV2 | |
| **-pAAV-Ng-S36D** | pAAV | Ng-S36D | CaMKIIα0.4 | AAV2 | |
| **pAAV-Ng-I33Q** | pAAV | Ng-I33Q | CaMKIIα0.4 | AAV2 | |
| **pAAV-Ng-EGFP** | pAAV | Ng | CaMKIIα0.4 | AAV2 | EGFP |
| **pFΔ6\*** |  | AAV helper |  | AAV2 |  |
| **pRV1\*** |  | cap |  | AAV2 |  |
| **pH21\*** |  | rep-cap |  | AAV2 |  |

\* Kindly donated by Dr Hilmar Bading (Department of Neurobiology, Interdisciplinary Center for Neurosciences (IZN), Heidelberg, Germany)
\*\* pAAV backbones were from Addgene #165429

**Supplementary Table 2.** Plasmids obtained from Addgene.

| Plasmids | Addgene | Backbone | Insert | promoter | expression | tag |
| --- | --- | --- | --- | --- | --- | --- |
| **pAAV-shNg-mRuby2** | Modified from #92155 | pAAV | shRNA targeting Ng | H1 | AAV2 | mRuby2 |
| **pAAV-shCTRL** | #181875 | pAAV | control shRNA | U6 | AAV2 | mCherry |
| **pCMVRδ8.74** | #22036 |  | gag pol tat rev |  | lentiviral |  |
| **pMD2.G** | #12259 |  | VSV-G envelope |  | lentiviral |  |

**Supplementary Table 3.** Antibodies used in this study for WB and Ifs.

| Antibodies | Species | Antibody dilution |  | Reference |
| --- | --- | --- | --- | --- |
|  |  | IF (cell culture) | WB |  |
| **Ankiryne-G** | Guinea-pig | 1:250 |  | Synaptic System, 386 005 |
| **βIII-tubulin** | Mouse |  | 1:50.000 | Sigma, T8660 |
| **Bax** | Rabbit |  | 1:2.000 | Santa Cruz Biotech., sc-493 |
| **Bcl-2** | Rabbit |  | 1:2.000 | Santa Cruz Biotech. sc-492 |
| **CaM** | Mouse |  | 1:5.000 | Millipore, 05-173 |
| **CaMKIIα** | Mouse |  | 1:10.000 | Millipore, 05-532 6G9 |
| **CaN-A** | Rabbit |  | 1:10.000 | Cell Signaling, 2614 |
| **Active caspase-3 (Asp175)** | Rabbit |  | 1:1.000 | Cell Signaling, 9961 |
| **eIF2α** | Rabbit |  | 1:2.000 | Santa Cruz Biotech. sc-133132 |
| **Phospho-CaMKII (Thr286)** | Rabbit |  | 1:2.000 | PhosphoSolutions, p1005-286 |
| **Phospho-eIF2 (Ser52)** | Rabbit |  | 1:2.000 | Invitrogen, 44-788G |
| **GAD65** | Mouse | 1:5.000 |  | DSHB Iowa GAD-6 |
| **GAPDH** | Mouse |  | 1:30.000 | Millipore, MAB374 6C5 |
| **Gephyrin-Oyster-550** | Mouse | 1:500 |  | Synaptic Systems, 147 011 C3 |
| **GluA1** | Rabbit |  | 1:2.000 | Millipore, AB1504 |
| **GluA2** | Rabbit |  | 1:2.000 | Cell Signaling #13607 |
| **GluN1** | Mouse |  | 1:2.000 | Millipore, MAB1586 |
| **GluN2A** | Rabbit |  | 1:2.000 | Millipore, AB1555P |
| **GluN2B** | Mouse |  | 1:2.000 | Millipore, MAB52220 |
| **MAP2** | Guinea pig<br>Rabbit<br>Chicken | 1:5000<br>1:2.000<br>1:1.000 |  | Synaptic Systems, 188004<br>Millipore AB5622<br>Abcam, ab5392 |
| **mGluR5** | Rabbit |  | 1:500 | Sigma-Aldrich, AB5675 |
| **Ng** | Rabbit | 1:1000 | 1:30.000 | Millipore, AB5620 |
| **PSD-95** | Mouse | 1:1000 |  | Millipore, MAB1596 clone 6G6 |
| **vGluT1** | Guinea-pig | 1:15.000 |  | Millipore, AB5905 |

## References

1. Represa A, Deloulme JC, Sensenbrenner M, et al (1990) Neurogranin: Immunocytochemical localization of a brain-specific protein kinase C substrate. J Neurosci 10:3782–3792. https://doi.org/10/gnfz34

2. Alvarez-Bolado G, Rodríguez-Sánchez P, Tejero-Díez P, et al (1996) Neurogranin in the development of the rat telencephalon. Neuroscience 73:565–580. https://doi.org/10/ds76rj

3. Watson JB, Sutcliffe JG, Fisher RS (1992) Localization of the protein kinase C phosphorylation/calmodulin-binding substrate RC3 in dendritic spines of neostriatal neurons. Proc Natl Acad Sci 89:8581–8585. 10.1073/pnas.89.18.8581

4. Gerendasy DD, Herron SR, Watson JB, Sutcliffe JG (1994) Mutational and biophysical studies suggest RC3/neurogranin regulates calmodulin availability. J Biol Chem 269:22420–22426

5. Huang KP, Huang FL, Jäger T, et al (2004) Neurogranin/RC3 enhances long-term potentiation and learning by promoting calcium-mediated signaling. J Neurosci 24:10660–10669. https://doi.org/10/dp9mnp

6. Baudier J, Bronner C, Kligman D, Cole RD (1989) Protein kinase C substrates from bovine brain. Purification and characterization of neuromodulin, a neuron-specific calmodulin-binding protein. J Biol Chem 264:1824–1828. https://doi.org/10/gnfz8s

7. Baudier J, Deloulme JC, Van Dorsselaer A, et al (1991) Purification and characterization of a brain-specific protein kinase C substrate, neurogranin (p17). Identification of a consensus amino acid sequence between neurogranin and neuromodulin (GAP43) that corresponds to the protein kinase C phosphorylation si. J Biol Chem 266:229–237

8. Hoffman L, Chandrasekar A, Wang X, et al (2014) Neurogranin Alters the Structure and Calcium Binding Properties of Calmodulin. J Biol Chem 289:14644–14655. https://doi.org/10/f56kv7

9. Casaletto KB, Elahi FM, Bettcher BM, et al (2017) Neurogranin, a synaptic protein, is associated with memory independent of Alzheimer biomarkers. Neurology 89:1782–1788. https://doi.org/10/gcptnh

10. Portelius E, Zetterberg H, Skillbäck T, et al (2015) Cerebrospinal fluid neurogranin: Relation to cognition and neurodegeneration in Alzheimer’s disease. Brain 138:3373–3385. https://doi.org/10/f3ncmw

11. Thorsell A, Bjerke M, Gobom J, et al (2010) Neurogranin in cerebrospinal fluid as a marker of synaptic degeneration in Alzheimer’s disease. Brain Res 1362:13–22. 10.1016/j.brainres.2010.09.073

12. Kester MI, Teunissen CE, Crimmins DL, et al (2015) Neurogranin as a Cerebrospinal Fluid Biomarker for Synaptic Loss in Symptomatic Alzheimer Disease. JAMA Neurol 72:1275–1280. 10.1001/jamaneurol.2015.1867

13. Kvartsberg H, Duits FH, Ingelsson M, et al (2015) Cerebrospinal fluid levels of the synaptic protein neurogranin correlates with cognitive decline in prodromal Alzheimer’s disease. Alzheimers Dement 11:1180–1190. https://doi.org/10/f2zkhv

14. Saunders T, Gunn C, Blennow K, et al (2023) Neurogranin in Alzheimer’s disease and ageing: A human post-mortem study. Neurobiol Dis 177:105991. 10.1016/j.nbd.2023.105991

15. Tzioras M, McGeachan RI, Durrant CS, Spires-Jones TL (2023) Synaptic degeneration in Alzheimer disease. Nat Rev Neurol 19:19–38. 10.1038/s41582-022-00749-z

16. Kvartsberg H, Lashley T, Murray CE, et al (2019) The intact postsynaptic protein neurogranin is reduced in brain tissue from patients with familial and sporadic Alzheimer’s disease. Acta Neuropathol (Berl) 137:89–102. https://doi.org/10/gd8x7g

17. Mons N, Enderlin V, Jaffard R, Higueret P (2001) Selective age-related changes in the PKC-sensitive, calmodulin-binding protein, neurogranin, in the mouse brain. J Neurochem 79:859–67. https://doi.org/10/fnt3ff

18. Iñiguez MA, De Lecea L, Guadano-Ferraz A, et al (1996) Cell-specific effects of thyroid hormone on RC3/neurogranin expression in rat brain. Endocrinology 137:1032–1041. 10.1210/endo.137.3.8603571

19. Zoeller RT, Bansal R, Parris C (2005) Bisphenol-A, an environmental contaminant that acts as a thyroid hormone receptor antagonist in vitro, increases serum thyroxine, and alters RC3/neurogranin expression in the developing rat brain. Endocrinology 146:607–612. 10.1210/en.2004-1018

20. Pak JH, Huang FL, Li J, et al (2000) Involvement of neurogranin in the modulation of calcium/calmodulin-dependent protein kinase II, synaptic plasticity, and spatial learning: A study with knockout mice. Proc Natl Acad Sci U S A 97:11232–11237. https://doi.org/10/dt8vbk

21. Krucker T, Siggins GR, McNamara RK, et al (2002) Targeted disruption of RC3 reveals a calmodulin-based mechanism for regulating metaplasticity in the hippocampus. J Neurosci 22:5525–5535. https://doi.org/10/gnf2pt

22. Zhabotinsky AM, Camp RN, Epstein IR, Lisman JE (2006) Role of the neurogranin concentrated in spines in the induction of long-term potentiation. J Neurosci 26:7337–7347. https://doi.org/10/b866m6

23. Kubota Y, Putkey JA, Waxham MN (2007) Neurogranin Controls the Spatiotemporal Pattern of Postsynaptic Ca2+/CaM Signaling. Biophys J 93:3848–3859. 10.1529/biophysj.107.106849

24. Ordyan M, Bartol T, Kennedy M, et al (2020) Interactions between calmodulin and neurogranin govern the dynamics of CaMKII as a leaky integrator. PLOS Comput Biol 16:e1008015. 10.1371/journal.pcbi.1008015

25. Miyakawa T, Yared E, Pak JH, et al (2001) Neurogranin null mutant mice display performance deficits on spatial learning tasks with anxiety related components. Hippocampus 11:763–775. https://doi.org/10/ch7jzf

26. Zhong L, Cherry T, Bies CE, et al (2009) Neurogranin enhances synaptic strength through its interaction with calmodulin. EMBO J 28:3027–3039. https://doi.org/10/bvhtj7

27. Petersen A, Gerges NZ (2015) Neurogranin regulates CaM dynamics at dendritic spines. Sci Rep 5:11135. https://doi.org/10/gnf2dm

28. Domínguez-González I, Vázquez-Cuesta SN, Algaba A, Díez-Guerra FJ (2007) Neurogranin binds to phosphatidic acid and associates to cellular membranes. Biochem J 404:31–43. https://doi.org/10/cr4f6x

29. Slemmon JR, Feng B, Erhardt JA (2000) Small proteins that modulate calmodulin-dependent signal transduction. Mol Neurobiol 22:99–113. 10.1385/MN:22:1-3:099

30. Randall Slemmon J, Morgan JI, Fullerton SM, et al (1996) Camstatins are peptide antagonists of calmodulin based upon a conserved structural motif in PEP-19, neurogranin, and neuromodulin. J Biol Chem 271:15911–15917. https://doi.org/10/b24x4b

31. Garrido-García A, de Andrés R, Jiménez-Pompa A, et al (2019) Neurogranin Expression Is Regulated by Synaptic Activity and Promotes Synaptogenesis in Cultured Hippocampal Neurons. Mol Neurobiol 56:7321–7337. 10.1007/s12035-019-1593-3

32. Kaech S, Banker G (2006) Culturing hippocampal neurons. Nat Protoc 1:2406–2415. 10.1038/nprot.2006.356

33. Gascón S, Paez-Gomez JA, Díaz-Guerra M, et al (2008) Dual-promoter lentiviral vectors for constitutive and regulated gene expression in neurons. J Neurosci Methods 168:104–112. https://doi.org/10/fkn5f7

34. McClure C, Cole KLH, Wulff P, et al (2011) Production and Titering of Recombinant Adeno-associated Viral Vectors. J Vis Exp JoVE e3348. 10.3791/3348

35. Schindelin J, Arganda-Carreras I, Frise E, et al (2012) Fiji: An open-source platform for biological-image analysis. Nat Methods 9:676–682. https://doi.org/10/f34d7c

36. Schneider CA, Rasband WS, Eliceiri KW (2012) NIH Image to ImageJ: 25 years of image analysis. Nat Methods 9:671–675. https://doi.org/10/gcwb4q

37. Gallego-Garcia C, Martínez Blanco E, Diez-Guerra FJ (2025) SynapTrack: An Automated Tool for Synapse Quantification

38. Grubb MS, Burrone J (2010) Activity-dependent relocation of the axon initial segment fine-tunes neuronal excitability. Nature 465:1070–1074. https://doi.org/10/fn7qnz

39. Zhang Y, Rózsa M, Liang Y, et al (2023) Fast and sensitive GCaMP calcium indicators for imaging neural populations. Nature 615:884–891. 10.1038/s41586-023-05828-9

40. Shaner NC, Campbell RE, Steinbach PA, et al (2004) Improved monomeric red, orange and yellow fluorescent proteins derived from Discosoma sp. red fluorescent protein. Nat Biotechnol 22:1567– 1572. https://doi.org/10/fw6mvd

41. Siebler M, Köller H, Stichel CC, et al (1993) Spontaneous activity and recurrent inhibition in cultured hippocampal networks. Synapse 14:206–213. 10.1002/syn.890140304

42. Charlesworth P, Cotterill E, Morton A, et al (2015) Quantitative differences in developmental profiles of spontaneous activity in cortical and hippocampal cultures. Neural Develop 10:1. 10.1186/s13064-014-0028-0

43. Martzen MR, Slemmon JR (1995) The Dendritic Peptide Neurogranin Can Regulate a Calmodulin-Dependent Target. J Neurochem 64:92–100. https://doi.org/10/cnnkwk

44. Bayer KU, Giese KP (2024) A revised view of the role of CaMKII in learning and memory. Nat Neurosci 1–11. 10.1038/s41593-024-01809-x

45. McIlwain DR, Berger T, Mak TW (2013) Caspase Functions in Cell Death and Disease. Cold Spring Harb Perspect Biol 5:a008656–a008656. https://doi.org/10/gbcrk4

46. D’Amelio M, Cavallucci V, Cecconi F (2010) Neuronal caspase-3 signaling: Not only cell death. Cell Death Differ 17:1104–1114. https://doi.org/10/ct5p46

47. Li Z, Jo J, Jia JM, et al (2010) Caspase-3 activation via mitochondria is required for long-term depression and AMPA receptor internalization. Cell 141:859–871. https://doi.org/10/b95r93

48. Ertürk A, Wang Y, Sheng M (2014) Local pruning of dendrites and spines by caspase-3-dependent and proteasome-limited mechanisms. J Neurosci 34:1672–1688. https://doi.org/10/f5r89b

49. Hollville E, Deshmukh M (2018) Physiological functions of non-apoptotic caspase activity in the nervous system. Semin Cell Dev Biol 82:127–136. 10.1016/j.semcdb.2017.11.037

50. Nguyen TTM, Gillet G, Popgeorgiev N (2021) Caspases in the Developing Central Nervous System: Apoptosis and Beyond. Front Cell Dev Biol 9:1910. https://doi.org/10/gnzhwc

51. Sarić N, Hashimoto-Torii K, Jevtović-Todorović V, Ishibashi N (2022) Nonapoptotic caspases in neural development and in anesthesia-induced neurotoxicity. Trends Neurosci 45:446–458. 10.1016/j.tins.2022.03.007

52. Ledda F, Paratcha G (2017) Mechanisms regulating dendritic arbor patterning. Cell Mol Life Sci 74:4511–4537. 10.1007/s00018-017-2588-8

53. Lohmann C, Wong ROL (2005) Regulation of dendritic growth and plasticity by local and global calcium dynamics. Cell Calcium 37:403–409. 10.1016/j.ceca.2005.01.008

54. Wayman GA, Impey S, Marks D, et al (2006) Activity-Dependent Dendritic Arborization Mediated by CaM-Kinase I Activation and Enhanced CREB-Dependent Transcription of Wnt-2. Neuron 50:897–909. 10.1016/j.neuron.2006.05.008

55. Grabrucker A, Vaida B, Bockmann J, Boeckers TM (2009) Synaptogenesis of hippocampal neurons in primary cell culture. Cell Tissue Res 338:333–341. https://doi.org/10/bvqgh8

56. Ivenshitz M, Segal M (2010) Neuronal density determines network connectivity and spontaneous activity in cultured hippocampus. J Neurophysiol 104:1052–1060. https://doi.org/10/ffb9g8

57. Han K-S, Cooke SF, Xu W (2017) Experience-Dependent Equilibration of AMPAR-Mediated Synaptic Transmission during the Critical Period. Cell Rep 18:892–904. 10.1016/j.celrep.2016.12.084

58. Díez-Guerra FJ (2010) Neurogranin, a link between calcium/calmodulin and protein kinase C signaling in synaptic plasticity. IUBMB Life 62:597–606. https://doi.org/10/b6s2cb

59. Zhong L, Gerges NZ (2012) Neurogranin targets calmodulin and lowers the threshold for the induction of long-term potentiation. PLoS ONE 7:1–8. https://doi.org/10/f343tq

60. Li L, Lai M, Cole S, et al (2020) Neurogranin stimulates Ca2+/calmodulin-dependent kinase II by suppressing calcineurin activity at specific calcium spike frequencies. PLOS Comput Biol 16:e1006991. 10.1371/journal.pcbi.1006991

61. Fernandes D, Carvalho AL (2016) Mechanisms of homeostatic plasticity in the excitatory synapse. J Neurochem 139:973–996. 10.1111/jnc.13687

62. Gulyaeva NV (2003) Non-apoptotic Functions of Caspase-3 in Nervous Tissue. Biochem Mosc 68:1171– 1180. 10.1023/B:BIRY.0000009130.62944.35

63. Li Z, Sheng M (2012) Caspases in synaptic plasticity. Mol Brain 5:15. 10.1186/1756-6606-5-15

64. Hyman BT, Yuan J (2012) Apoptotic and non-apoptotic roles of caspases in neuronal physiology and pathophysiology. Nat Rev Neurosci 13:395–406. https://doi.org/10/f3zd9h

65. Mukherjee A, Williams DW (2017) More alive than dead: non-apoptotic roles for caspases in neuronal development, plasticity and disease. Cell Death Differ 24:1411–1421. 10.1038/cdd.2017.64

66. Jing G, Yuan K, Liang Q, et al (2012) Reduced CaM/FLIP binding by a single point mutation in c-FLIPL modulates Fas-mediated apoptosis and decreases tumorigenesis. Lab Invest 92:82–90. 10.1038/labinvest.2011.131

67. Li Z, Jo J, Jia J-M, et al (2010) Caspase-3 Activation via Mitochondria Is Required for Long-Term Depression and AMPA Receptor Internalization. Cell 141:859–871. 10.1016/j.cell.2010.03.053

68. Iñiguez MA, Rodriguez-Peña A, Ibarrola N, et al (1992) Adult rat brain is sensitive to thyroid hormone. Regulation of RC3/neurogranin mRNA. J Clin Invest 90:554–8. https://doi.org/10/bz8324

69. Stefansson H, Ophoff RA, Steinberg S, et al (2009) Common variants conferring risk of schizophrenia. Nature 460:744–7. https://doi.org/10/cjdkn5

70. Xue M, Sun F-R, Ou Y-N, et al (2020) Association of cerebrospinal fluid neurogranin levels with cognition and neurodegeneration in Alzheimer’s disease. Aging 12:9365–9379. 10.18632/aging.103211

